# Neuropeptide Y co-opts neuronal ensembles for memory lability and stability

**DOI:** 10.1101/2024.05.09.593455

**Authors:** Yan-Jiao Wu, Xue Gu, Yalei Kong, Shuo Yang, Huan Wang, Miao Xu, Qi Wang, Xin Yi, Ze-Jie Lin, Zhi-Han Jiao, Hoiyin Cheung, Xin-Yu Zhao, Xin Bian, Qin Jiang, Ying Li, Michael X. Zhu, Lu-Yang Wang, Yulong Li, Ju Huang, Qian Li, Wei-Guang Li, Tian-Le Xu

## Abstract

Memory engrams are formed by activity-dependent recruitment of distinct subsets of excitatory principal neurons (or neuronal ensembles) whereas inhibitory neurons pivot memory lability and stability^1–5^. However, the molecular logic for memory engrams to preferentially recruit specific type of interneurons over other subtypes remains enigmatic. Using activity-dependent single-cell transcriptomic profiling^6–8^ in mice with training of cued fear memory and extinction, we discovered that neuropeptide Y (NPY)-expressing (NPY^+^) GABAergic interneurons in the ventral hippocampal CA1 (vCA1) region exert fast GABAergic inhibition to facilitate the acquisition of memory, but bifurcate NPY-mediated slow peptidergic inhibition onto distinct sub-ensembles underlying the extinction of single memory trace. Genetically encoded calcium and NPY sensors revealed that both calcium dynamics of NPY^+^ neurons and their NPY release in vCA1 ramp up as extinction learning progresses while behavioral state switches from “fear-on” to “fear-off”. Bidirectional manipulations of NPY^+^ neurons or NPY itself demonstrated NPY is both necessary and sufficient to control the rate and degree of memory extinction by acting on two physically non-overlapping sub-ensembles composed of NPY1R- and NPY2R-expressing neurons. CRISPR/Cas9-mediated knockout of NPY2R or NPY1R further unravels that NPY co-opts its actions on these two sub-ensembles to gate early fast and late slow stages of extinction. These findings exemplify the intricate spatiotemporal orchestration of slow peptidergic inhibitions from single subtype of GABAergic interneurons to fine-tune engram lability verse stability of memory.

## Main

Growing number of studies have demonstrated that any given memory trace is formed by a distinct subset of cells being recruited to encode the experience during learning. Activity-dependent genetic tagging approaches showed that sparsely distributed neuronal ensembles are activated during specific learning epochs to cascade memory engrams in time and space^1–5^. Among these activity-defined engram neurons, the excitatory principal neurons with increased intrinsic excitability and excitatory synaptic plasticity are most likely recruited to construct engram-to-engram connectivity for memory expression in a dynamic manner^3,5,9^. Inhibition by GABAergic interneurons suppresses non-engram cells and maintains homeostasis of neural networks to specialize memory engrams^10,11^, gating convertibility for memory fate^12,13^. Recently, specific subtypes of GABAergic interneurons have emerged as the key components of memory engram networks^5,14^. GABAergic interneurons exhibit high heterogeneity in their location and projection to control spatiotemporal excitation of engram cells and some interneurons co-release peptides, such as somatostatin (SST), cholecystokinin (CCK), neuropeptide Y (NPY), and others. Although these peptides have been widely utilized as cell-specific molecular markers^15–17^, fundamental questions arise whether GABA, or peptides alone, or their co-release from different types of interneurons drive divergent or convergent behavioral outcomes. These uncertainties are further mystified by two major outstanding questions: Which types of inhibitory interneurons are recruited during memory expression and/or its extinction (a typical form of memory-based behavioral state switch)^18,19^? and under what circumstances are peptides co-released to regulate learning and memory? To address these knowledge gaps, we investigated the dynamic engagements of hippocampal GABAergic interneurons in instructing neuronal ensembles for memory expression and extinction.

### *Npy^+^* inhibitory neurons are selectively activated in ventral hippocampus during extinction learning

Fear memory^20^ and its extinction^19^ serve as paradigmatic examples illustrating the dynamic balance between memory lability and stability. These processes rely on the fluid organization of contrasting and interconvertible memory engrams, which are crucial for organisms to flexibly adapt to threatening environments. Dysregulated adaptions likely underlie anxiety and trauma-related disorders^21,22^. Fear conditioning establishes associations, known as fear memory, between threatening experiences (i.e., foot shock) and neutral stimuli (i.e., sound)^20^. In contrast, extinction training involves exposing the conditioned neutral stimuli without threatening reinforcement, decoupling the cue for associative memory by forming new extinction memory engram^19^. Successful extinction suppresses the original fear memory to a dormant state with the potential for spontaneous relapse over time^23–25^. Taking advantage of fear memory and extinction as robust platforms to differentiate specific neuronal ensembles and their interconvertibility at different stages, we first profiled transcriptomic changes at the single-cell level to identify molecular signatures associated with two memory engrams^8,26,27^. To this end, we used unbiased high-throughput droplet-based single-cell RNA sequencing (scRNA-seq) combined with a single-cell preparation method designed to eliminate transcriptional perturbations during sample preparation, referred to as Act-seq^6^. This method minimizes artificially induced transcriptional perturbations, enabling the faithful detection of acute changes in individual cells induced by behavioral activity.

We focused on the ventral hippocampal CA1 region (vCA1) because of its role as a crucial hub for fear extinction^28–32^. Wild-type mice underwent auditory fear conditioning on Day 1 when a foot shock was associated with a neutral auditory tone before being subjected to extinction training on Day 2 when 20-trial conditioned stimulus (CS) presentations were presented in a new context without foot shock (Ext group). As a control, mice were exposed to 2-trial CS presentations, inducing heightened fear responses (No Ext group). Two hours after behavioral tests, samples from the vCA1 of both No Ext and Ext mice were collected and dissociated into single-cell suspensions (Fig. 1a, b).

**Fig 1.**
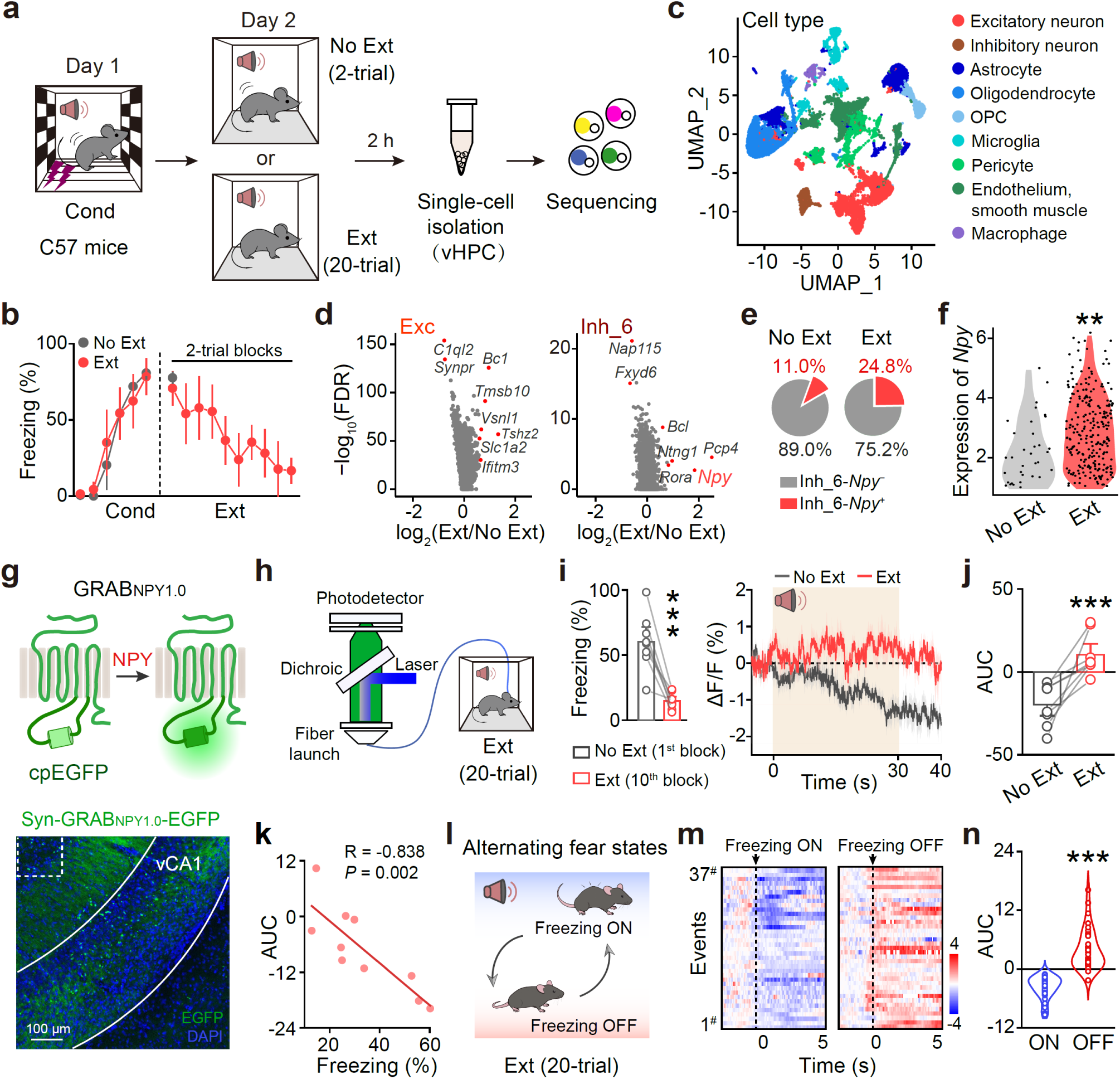
Dynamics of vCA1 extracellular NPY during extinction learning. **a**, Schematic illustrating the behavioral procedure for No Ext or Ext, and Act-Seq. **b**, Freezing levels of the Ext group were significantly lower compared to those of the No Ext group. n = 4 and 4 mice for No Ext and Ext, respectively. **c**, Uniform Manifold Approximation and Projection (UMAP) resolves 46,894 cells from vHPC into nine types. Colors denote main cell types. **d**, Sample volcano plots for inhibitory and excitatory neurons indicating genes identified as extinction stimulus-regulated (|log2 fold change| > 0.585 and FDR < 0.05) for No Ext and Ext comparisons. **e**, Pie charts demonstrate a higher proportion of *Npy*^+^ neurons among all inhibitory neurons in Ext mice compared to No Ext mice. **f**, Violin plots depict a notable increase in *Np*y expression levels among *Npy*^+^ neurons in the Ext group. \*\**P* = 0.001, unpaired Student’s *t*-test. Each dot represents a single cell. **g**, Schematic of the GRAB_NPY1.0_ sensor and a representative image showing the expression in vCA1. **h**, Schematic of fiber photometry recording during extinction learning. **i**, A significant increase in the response of GRAB_NPY1.0_ in the Ext state (the last two CS presentation, i.e., the 10^th^ blocks, with diminished fear response) compared to the No Ext state (the first two CS presentation, i.e., the 1^st^ blocks, with heighted fear response). n = 8 mice. Freezing levels, \*\*\**P* = 0.0004, two-tailed paired Student’s *t*-test. **j**, Area Under the Curve (AUC) of the ΔF/F GRAB_NPY1.0_ fluorescence during Ext and No Ext state, respectively. n = 8 mice. \*\**P* = 0.002, two-tailed paired Student’s *t*-test. **k**, Increased GRAB_NPY1.0_ sensor response showed a negative correlation with diminished freezing levels during extinction. Pearson Corr. = –0.838, \*\**P* = 0.002. **l**, Behavioral state alternated between freezing ON and OFF during extinction learning. **m, n**, Heatmaps (**m**) and AUC (**n**) of GRAB_NPY1.0_ fluorescence for 5 s following freezing ON and OFF epochs. n = 8 mice and 37 events for freezing ON and OFF, respectively. \*\*\**P* < 0.0001, unpaired Student’s *t*-test.

A total of 46,894 individual cells, including 20,263 from No Ext and 26,631 cells from Ext groups of mice, were profiled by uniform manifold approximation and projection (UMAP) for dimensionality reduction. Unsupervised clustering identified 35 distinct clusters across all samples (Fig. 1c), manually identified based on the expression of known cell-type-specific markers (Extended Data Fig. 1a). Various cell clusters exhibited differential responsiveness to Ext over No Ext treatment (Extended Data Fig. 1b). These changes diverged to induce different expression levels of immediate early genes (IEGs) across different cell types and clusters (Extended Data Fig. 2). Among the neuron types, Ext caused upregulation of 6 genes in excitatory neurons and 5 genes in inhibitory neurons (Fig. 1d). In excitatory neurons, Ext triggered varied transcriptomic program changes in all the 8 clusters (Extended Data Fig. 3a–h). The expression level of *Fos*, classically considered as a hallmark of active neuronal ensembles responding to salient stimuli^25,33,34^, was significantly increased in Clusters 10 and 32 excitatory neurons of the Ext compared to the No Ext group (Extended Data Fig. 3i). Inhibitory neurons from Cluster 6 upregulated the expression of *Pcp4* and *Npy* genes, up to 2-fold by the Ext (Fig. 1d).

We then conducted dual-color fluorescence in situ hybridization (dFISH) analyses to corroborate the Act-seq results. The expression of *Npy* was exclusive to VGAT^+^ (vesicular GABA transporter-expressing) neurons rather than Vglut1^+^ (vesicular glutamate transporter 1-expressing) neurons, confirming the selective expression of *Npy* in vCA1 GABAergic inhibitory interneurons. Additionally, the *Npy* signal was more enchained in the stratum oriens (SO) of CA1 region^35^ (Extended Data Fig. 4). Further analysis of Act-seq results showed a higher proportion of *Npy*^+^ neurons among all inhibitory neurons^17^ in Ext mice compared to No Ext mice (Fig. 1e). Moreover, these *Npy*^+^ neurons exhibited a notable increase in *Npy* expression levels (Fig. 1f), that significantly correlated with the changes of IEGs within each inhibitory neuron in the Ext group, but not those in the No Ext group (Extended Data Fig. 5). Consistently, dFISH analyses showed a significant increase in the percentage of *Npy*^+^ and *Npy*^+^/*Fos*^+^ co-labeled neurons in the vCA1 of Ext compared to that of No Ext mice. Furthermore, the expression level of *Npy* and *Fos* in the vCA1 exhibited a significant correlation in the Ext group but not the No Ext group (Extended Data Fig. 6). Together, these analyses validated *Npy*^+^ GABAergic interneurons being transcriptionally activated in the vCA1 region during fear extinction can potentially serve as the key cell type to gate extinction learning.

### Dynamics of vCA1 extracellular NPY during extinction learning

NPY is a 36-amino acid peptide neurotransmitter implicated in physiological processes such as energy homeostasis^36^, learning and memory^37,38^, acting on different types of Gi/o-coupled NPY receptors^39^. NPY also plays a role in regulating memory systems relevant to pathophysiology of post-traumatic stress disorder (PTSD) with therapeutic potential^40^, yet its brain-region- and cell-type-specific origin, spatiotemporal targeting and signaling mechanisms in fear memory remain ambiguous. To directly monitor vCA1 NPY dynamics during fear extinction, we targeted expression of a genetically encoded fluorescent NPY sensor^41^ termed as GRAB_NPY1.0_ (Fig. 1g) to vCA1 neurons and quantify real-time NPY release *in vivo* during extinction learning using fiber photometry (Fig. 1h). We found a significant increase in CS-evoked NPY release in the Ext state (the last two CS presentation, i.e., the 10^th^ block, with diminished fear response) compared to the No Ext state (the first two CS presentation, i.e., the 1^st^ block, with heightened fear response) (Fig. 1i, j). Additionally, NPY release to the repeated CS presentation ramped up during extinction learning while fear responses diminished (Fig. 1k). Throughout the 30-s period of each CS trail during extinction learning, the behavioral state of mice alternated between freezing ON and freezing OFF epochs (Fig. 1l), with decrease and increase of the NPY signal in parallel (Fig. 1m, n). In contrast, neither a non-ligand binding mutant form of the sensor (GRAB_NPYmut_) in wild-type mice nor the GRAB_NPY1.0_ sensor in *Npy* knockout mice exhibited any detectable response in vCA1 during extinction learning (Extended Data Fig. 7).

### Dynamics of vCA1 NPY^+^ neuronal activity during extinction learning

To elucidate the relationship between extracellular NPY concentrations and NPY^+^ interneurons in the vCA1 and assess the extent to which these neurons contribute to NPY release associated with extinction learning, we expressed Cre-dependent, red-shifted opsin (ChrimsonR), along with the GRAB_NPY1.0_, into the vCA1 of NPY-Cre mice to monitor NPY sensor responses specifically upon activation of NPY^+^ interneurons (Fig. 2a). Our results confirmed that stimulating NPY^+^ interneurons was sufficient to evoke NPY release in the vCA1 (Fig. 2b, c). We further assessed the neuronal activity of vCA1 NPY^+^ neurons by expressing a Cre-dependent genetically encoded calcium indicator GCaMP6m in NPY-Cre mice and continuously recorded NPY^+^ neuronal calcium activity using fiber photometry (Fig. 2d). Calcium responses during the Ext state were significantly higher than those during the No Ext state (Fig. 2e, f). As extinction progressed, calcium responses in NPY^+^ neurons increased, showing a significant negative correlation with the decreased fear response during intermittent CS presentations (Fig. 2g). Consistent with the NPY sensor responses, we noted a decrease in calcium signals in NPY^+^ neurons during freezing ON and an increase when the mice were freezing OFF (Fig. 2h, i). These results thus suggested the active involvement of vCA1 NPY^+^ neuronal activity in the dynamics of NPY release during fear extinction, underscoring its potential role in driving progressive extinction.

**Fig 2.**
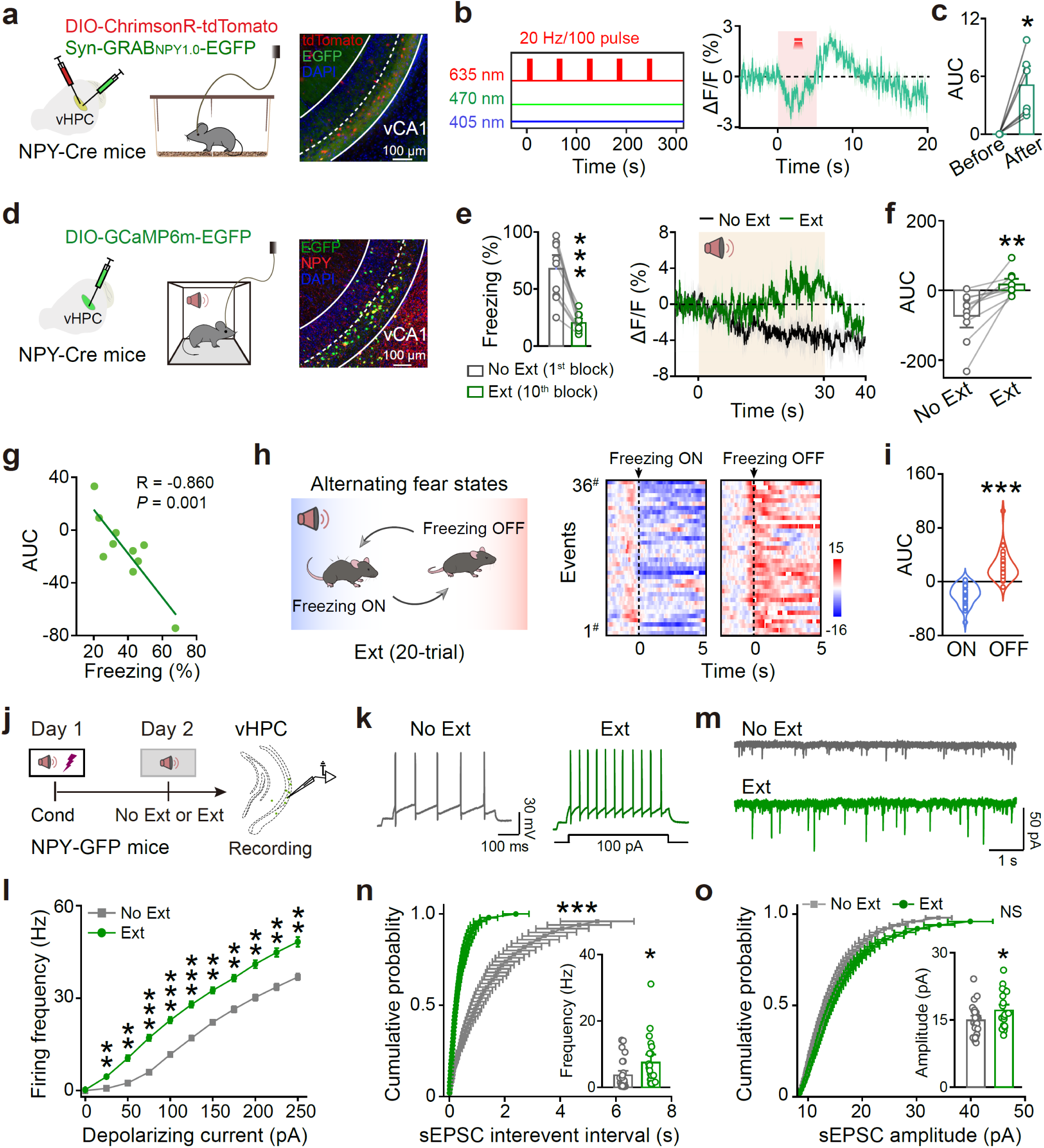
Dynamics of vCA1 NPY^+^ neuronal activity during extinction learning. **a**, Schematic of viral injection for fiber photometry and a representative image showing the expression of ChrimsonR and GRAB_NPY1.0_ sensors in vCA1. **b**, Averaged trace of the change in GRAB_NPY1.0_ fluorescence in response to stimulating NPY^+^ interneurons with 100 pulses at 20 Hz frequency. **c**, AUC of the ΔF/F GRAB_NPY1.0_ sensors fluorescence for 5 s before and after NPY^+^ neuronal activation, respectively. n = 6 mice. \**P* = 0.012, two-tailed paired Student’s *t*-test. **d**, Schematic of viral injection for fiber photometry and a representative image showing the overlap of GCaMP6m and the labeled NPY^+^ populations in vCA1. **e**, A significant increase in the response of GCaMP6m in the Ext state (the last two CS presentation, i.e., the 10^th^ blocks, with diminished fear response) compared to the No Ext state (the first two CS presentation, i.e., the 1^st^ blocks, with heighted fear response) was increased during the Ext state compared to the No Ext state. Freezing levels, \*\*\**P* < 0.0001, two-tailed paired Student’s *t*-test. **f**, AUC of the ΔF/F GCaMP6m fluorescence during Ext and No Ext state, respectively. n = 10 mice. \*\**P* = 0.002, two-tailed paired Student’s *t*-test. **g**, During extinction, increased calcium responses in NPY^+^ neurons showed a significant negative correlation with the decreased fear response. Pearson Corr. = –0.860, \*\**P* = 0.001. **h, i**, Heatmaps (**h**) and AUC (**i**) of NPY^+^ neuronal calcium responses for 5 s following freezing ON and OFF epochs. n = 10 mice and 36 events for freezing ON and OFF, respectively. \*\*\**P* < 0.0001, unpaired Student’s *t*-test. **j**, Schematic showing behavioral procedure for electrophysiological recordings of NPY^+^ interneurons in vHPC acute slices. **k, m**, Representative traces showing voltage responses to +100 pA current injections (**k**) and spontaneous excitatory postsynaptic current (sEPSC) (**m**) of NPY^+^ interneurons. **l**, The frequency of action potential (AP) discharge as a function of step-current intensity (0–250 pA, 500 ms). No Ext, n = 26 neurons of three mice; Ext, n = 31 neurons of three mice. \*\**P* < 0.01, \*\*\**P* < 0.001, unpaired Student’s *t*-test. **n**, Cumulative probability of the sEPSC inter-event intervals. \*\*\**P* < 0.0001, two-sample Kolmogorov-Smirnov test. Bar graphs for amplitude, \**P* = 0.035, unpaired Student’s *t*-test. **o**, Cumulative probability of the sEPSC amplitude. NS (not significant), *P* = 0.967, two-sample Kolmogorov-Smirnov test. Bar graphs for frequency, \**P* = 0.045, unpaired Student’s *t*-test.

We next conducted a comparative analysis of the excitability of NPY^+^ interneurons between the No Ext and Ext groups using patch-clamp electrophysiology. A substantial increase in the frequency of action potentials evoked by the same current step injections in the Ext group was observed, while the amplitude remained unchanged (Fig. 2j–l). Furthermore, the frequency of spontaneous excitatory postsynaptic currents (sEPSC) as indicators of global excitatory synaptic inputs to NPY^+^ interneurons showed a significant rise in the Ext group, with less pronounced changes in amplitude (Fig. 2m–o). These findings elucidate that throughout the process of extinction learning, successive exposures to the CS incrementally enhance excitatory drive to NPY^+^ interneurons as well as their - excitability in vCA1, ramping up the activity of NPY^+^ neurons to release NPY. This, in turn, contributes to the suppression of fear memory and the emergence of extinction memory. These data suggested that during extinction learning, activated NPY^+^ GABAergic interneurons in the vCA1, with endogenous release of NPY, predict the dynamics of fear and extinction memory.

### Manipulation of vCA1 NPY^+^ interneuron activity controls memory extinction

To evaluate the necessity and efficacy of NPY^+^ neuronal activity in extinction learning, we introduced halorhodopsin (NpHR) or channelrhodopsin-2 (ChR2) into NPY^+^ interneurons in the vCA1 and manipulated their activity using spatially targeted light (Fig. 3a–c). Optogenetic inhibition of NPY^+^ interneurons during CS presentation in extinction learning intensified the tone-shock association, sustaining a higher fear response (Fig. 3d). Even during extinction retrieval without optogenetic inhibition, the NpHR group maintained a higher level of conditioned fear. Conversely, optogenetic activation of NPY^+^ neurons during CS presentation in extinction learning disassociated the tone-shock association, resulting in reduced fear responses both in extinction learning and retrieval (Fig. 3e). Further analyses of extinction time course showed two stages of decline in cued freezing levels over 10-block epochs (20-trail CS): Stage 1, i.e., the first 5 blocks, being rapid, likely reflects the lability of fear memory to extinction; and Stage 2, the last 5 blocks, being much slower, likely measures the stability of fear memory (Fig. 3f and Extended Data Fig. 8a–d). Optogenetic inhibition of NPY^+^ interneurons largely ablated Stage 1 extinction and precluded its progression to Stage 2; and conversely activation of NPY^+^ neurons robustly augmented the onset of Stage 1 extinction but had little effect on the stable setpoint of freezing level for Stage 2 (Fig. 3f and Extended Data Fig. 8a–d). These observations indicated that Stage 1 and 2 of extinction are interlocked processes, for which the activity level of NPY^+^ interneurons may confer sequentially labile and stable states of extinction memory.

**Fig 3.**
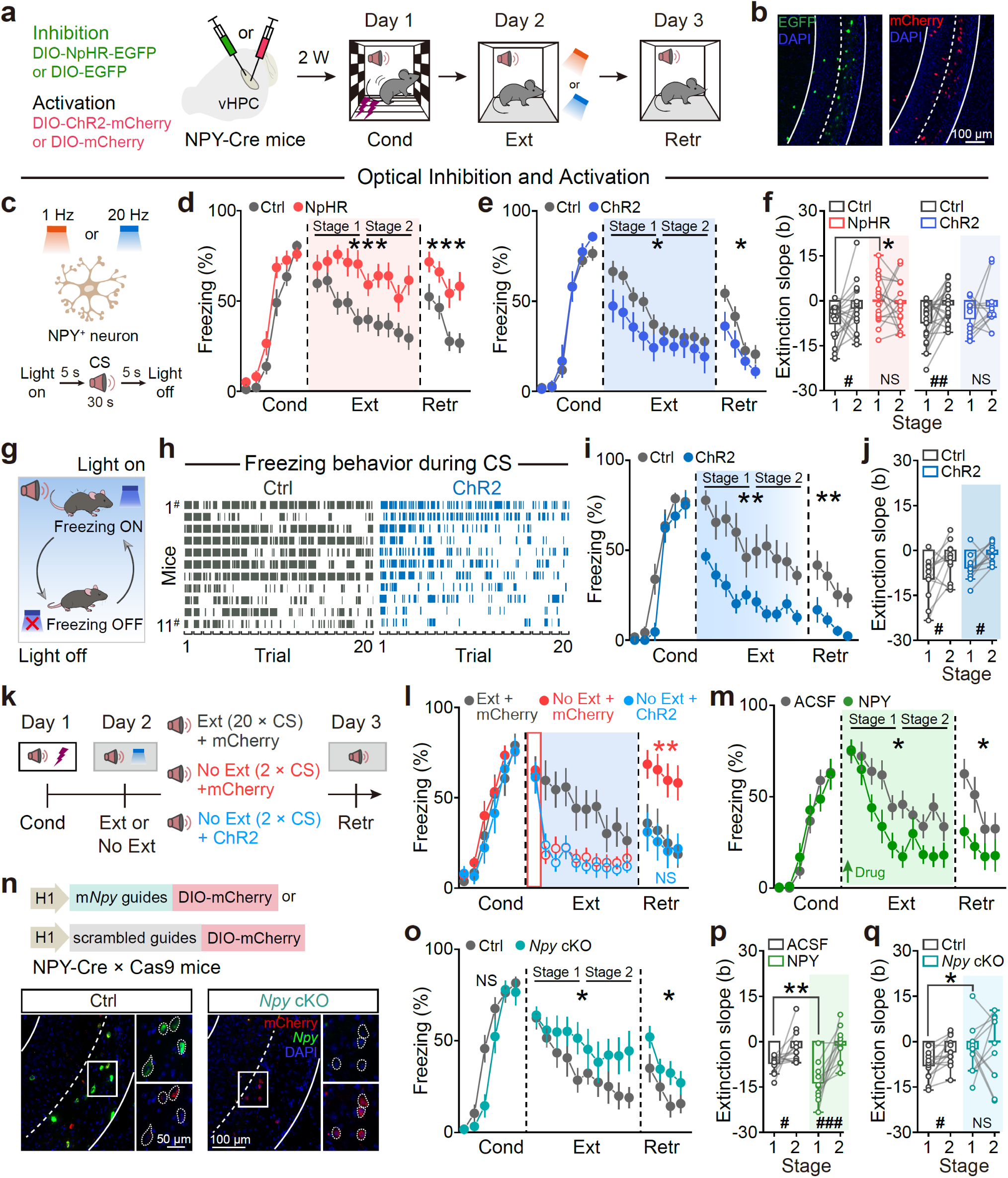
Manipulation of both vCA1 NPY^+^ interneurons activity and NPY dynamics controls memory extinction. **a**, Schematics of viral injection and behavioral procedure for optogenetic manipulations of NPY^+^ neuronal activity in the vCA1. **b**, Representative images of NpHR-EGFP and ChR2-mCherry expression in NPY^+^ neurons. **c**, Photostimulation protocol was performed during extinction. **d**, Inhibition of NPY^+^ neurons during extinction impeded extinction learning. n = 20 mice for Ctrl and 16 mice for NpHR. Two-way repeated ANOVA, main effects of group, Ext: F_(1, 34)_ = 19.036, \*\*\**P* = 0.0001. Retr: F_(1, 34)_ = 12.030, \*\**P* = 0.001. **e**, Activation of NPY^+^ neurons during extinction facilitated extinction memory. n = 22 mice for Ctrl and 12 mice for ChR2. Two-way repeated ANOVA, main effects of group. Ext: F_(1, 32)_ = 5.38, \**P* = 0.027. Retr: F_(1, 32)_ = 4.770, \**P* = 0.036. **f**, Through linear fitting for extinction Stage 1 and Stage 2, the extinction slope during Stage 1 was significantly greater than that during Stage 2 in the control group, but not optogenetic stimulation group. Optogenetic inhibition of NPY^+^ neurons largely ablated Stage 1 extinction. Stage 1 *vs.* Stage 2: Ctrl for NpHR, ^#^*P* = 0.030; NpHR, NS, *P* = 0.719; Ctrl for ChR2, ^##^*P* = 0.006, ChR2, NS, *P* = 0.189, two-tailed paired Student’s *t*-test. Ctrl *vs.* NpHR, \*\**P* = 0.002, unpaired Student’s *t*-test. **g**, Schematics of the behavioral procedure for optogenetic activation of NPY^+^ neurons when the mice exhibited freezing throughout the 30-s period of each CS trail in extinction. **h**, Freezing levels corresponds to optogenetic activation patterns in Ctrl and ChR2 groups. **i**, Activating NPY^+^ neurons during the CS-induced freezing ON epoch significantly facilitated fear extinction. n = 11 mice for Ctrl, and 11 mice for ChR2. Two-way repeated ANOVA, main effects of group. Ext: F_(1, 20)_ = 15.769, \*\**P* = 0.001. Retr: F_(1, 20)_ = 14.604, \*\**P* = 0.001. **j**, Optogenetic activation of NPY^+^ neurons during the CS-induced freezing ON epoch did not alter the dynamics of extinction learning. Stage 1 *vs.* Stage 2: Ctrl, ^#^*P* = 0.044; ChR2, ^#^*P* = 0.012, two-tailed paired Student’s *t*-test. **k**, Schematics for behavioral procedure to activate NPY^+^ neurons following fear retrieval (2 × CS exposure) but omitted the subsequent 18 × CS exposure for extinction training. **l**, Activation of NPY^+^ neurons without the entire learning procedure achieved similar fear reduction effects as the control extinction group. n = 10 mice for Ext + mCherry, n = 14 mice for No Ext + mCherry, n = 11 mice for No Ext + ChR2. Two-way repeated ANOVA, main effects of group. Ext: F_(2, 32)_ = 7.487, \*\**P* = 0.002; Ext + mCherry *vs.* No Ext + mCherry, \*\**P* = 0.002; Ext + mCherry *vs.* No Ext + ChR2, \*\**P* = 0.001; No Ext + mCherry *vs.* No Ext + ChR2, NS, *P* = 0.690. Retr: F_(2, 32)_ = 8.020, \*\**P* = 0.002; Ext + mCherry *vs.* No Ext + mCherry, \*\**P* = 0.003; Ext + mCherry *vs.* No Ext + ChR2, NS, *P* = 0.790; No Ext + mCherry *vs.* No Ext + ChR2, \*\**P* = 0.001. **m, p**, Microinjection of NPY into vHPC increased the rate and degree of fear extinction. n = 12 mice for ACSF, and 11 mice for NPY. (**m**) Two-way repeated ANOVA, main effects of group. Ext: F_(1, 21)_ = 6.189, \**P* = 0.021. Retr: F_(1, 21)_ = 5.186, \**P* = 0.033. (**p**) Stage 1 *vs.* Stage 2: ACSF, ^#^*P* = 0.039; NPY, ^###^*P* = 0.0008, two-tailed paired Student’s *t*-test. ACSF *vs.* NPY, \*\**P* = 0.004, unpaired Student’s *t*-test. **n**, Schematic of *Npy* gene conditional knockout (cKO) in the vCA1 NPY^+^ neurons and images showing the expression of *Npy* and mCherry in the control and *Npy* cKO groups. **o, q**, *Npy* cKO in vCA1 NPY^+^ populations significantly impeded extinction and eliminated the difference of extinction slope between the first and second halves during extinction learning. n = 13 mice for Ctrl, and 12 mice for *Npy* cKO. (**o**) Two-way repeated ANOVA, main effects of group. Ext: F_(1, 23)_ = 4.572, \**P* = 0.043. Retr: F_(1, 23)_ = 4.903, \**P* = 0.037. (**q**) Stage 1 *vs.* Stage 2: Ctrl, ^#^*P* = 0.011; *Npy* cKO, NS, *P* = 0.446, two-tailed paired Student’s *t*-test. Ctrl *vs. Npy* cKO, \**P* = 0.038, unpaired Student’s *t*-test.

To reinforce the proposed roles of NPY^+^ neuronal activity in regulating the liability and stability of fear extinction (Fig. 3g–j and Extended Data Fig. 8e, f), we repeated the optogenetic activation experiment following fear retrieval (2 × CS exposure) but omitted the subsequent 18 × CS exposure for extinction training (Fig. 3k). Activation of NPY^+^ neurons without the entire learning procedure accelerated Stage 1 extinction achieved similar fear reduction effects as the control extinction group (Fig. 3l). The effects of endogenous NPY levels appeared to be specific to fear extinction because locomotion of these mice is not altered by the same manipulations (Extended Data Fig. 9). These findings suggested that manipulating vCA1 NPY^+^ interneurons bidirectionally dictate and even instate extinction, with their action being more prominent during the behavioral state switch from “fear-on” to “fear-off” during extinction learning.

### NPY itself is both necessary and sufficient for fear extinction

To investigate whether the NPY peptide itself played a similar role as vCA1 NPY^+^ neuron activation in fear extinction, we microinjected exogenous NPY into the vCA1 through an implanted cannula 15 min prior to extinction leaning. We observed significant increases in the rate and degree of fear extinction (Fig. 3m, p), phenocopying the effects of activating NPY^+^ interneurons (Fig. 3e).

Conversely, using *Npy* gene knockout (*Npy^−/−^*) or knockdown (*Npy^+/−^*) mice, where the expression of NPY was undetectable or significantly downregulated, we observed impaired fear extinction, without affecting fear conditioning, and the difference in extinction slope between Stage 1 and 2 during extinction training was lost (Extended Data Fig. 10). For the location-specific genetic deletion of *Npy* in vCA1, we disrupted *Npy* expression from vCA1 NPY^+^ neurons while preserving their GABAergic machinery through Cre-dependent CRISPR-Streptococcus pyogenes Cas9 (SpCas9) with guide RNAs being targeted to exons 2 and 3 of the *Npy* gene in NPY-Cre::Cas9 (lox-stop-lox-Cas9) double transgenic mice. The Cas9 protein was specifically expressed in NPY^+^ neurons and induced a double-strand break in the *Npy* gene targeted by the *Npy* guide RNAs, leading to the formation of loss-of-function insertion-deletion (indel) mutations by non-homologous end joining, enabling site-specific targeted mutation of the *Npy* gene *in vivo*. We confirmed that the adeno associated virus (AAV) carrying such *Npy* guide RNAs effectively reduced *Npy* gene expression in NPY^+^ neurons compared to the control AAV with a scramble sequence (Fig. 3n). The selective knockout of *Npy* in vCA1 NPY^+^ populations significantly impeded extinction but did not affect fear conditioning (Fig. 3o) and eliminated the difference of extinction slope between the first and second halves during extinction learning (Fig. 3q and Extended Data Fig. 11). Thus, in line with the optogenetic manipulation of vCA1 NPY^+^ neuronal activity, changing the NPY peptide levels in this region bidirectionally influences the efficacy of fear extinction.

Given that NPY^+^ neurons are GABAergic interneurons in the vCA1 (Extended Data Fig. 4), we investigated whether the co-released GABA by NPY^+^ neurons is involved in fear and extinction regulation. We expressed Cre-dependent ChrimsonR along with the GRAB_GABA0.8_ sensor in the vCA1 of NPY-Cre mice to monitor GABA sensor responses specifically upon activation of NPY^+^ neurons (Extended Data Fig. 12a). Our results showed that artificial stimulation of NPY^+^ neurons in the vCA1 induced the release of NPY (Fig. 2a–c) accompanied by GABA (Extended Data Fig. 12a, b). When we inactivated GABA release from vCA1 NPY^+^ neurons while preserving NPY expression through Cre-dependent CRISPR-SpCas9 by targeting exons 2 and 3 of the *slc32a1* gene in NPY-Cre::Cas9 double transgenic mice, we found that selective ablation of GABA release from vCA1 NPY^+^ neurons prevented fear conditioning (Extended Data Fig. 12c, d). Furthermore, inhibition of NPY and GABA co-release by optogenetic inhibition of NPY^+^ neuronal activity during conditioning, resulted in decreased fear levels during retrieval (Extended Data Fig. 12e, f). This finding indicated distinctive roles of GABA and NPY in memory process of hippocampus (Fig. 3n, o and Extended Data Fig. 10c). NPY^+^ neurons in other brain regions are unlikely involved because manipulations of NPY^+^ neurons in the hypothalamic arcuate nucleus (ARC) had no impact on fear memory (Extended Data Fig. 13). Fast GABA release by NPY^+^ interneurons plays an indispensable role in memory encoding, while slow NPY release primarily mediates memory extinction locally in hippocampus.

### Non-overlapping neurons with distinct NPY receptor subtypes

To further unravel the downstream targets of NPY in fear extinction, we delved into single-cell transcriptomic data, revealing the expression of three primary types of NPY receptor genes — *Npy1r*, *Npy2r*, and *Npy5r* — in various cell types of the vCA1. Excitatory and inhibitory neurons predominantly exhibited higher expression levels of *Npy1r* and *Npy2r* compared to *Npy5r* (Extended Data Fig. 14a–f). Further analysis disclosed that *Npy1r* and *Npy2r* were essentially expressed in largely non-overlapping neuronal populations (Fig. 4a, b), as validated by dFISH analyses (Fig. 4c). Along the anatomic organization^42^ of hippocampal field CA1, *Npy1r* displayed specific cytoarchitectural distributions — from the superficial (vCA1-sps) to the middle (vCA1-spm) to the deep (vCA1-spd) pyramidal layers — gradually decreasing in expression (Fig. 4c, d and Extended Data Fig. 14g). Conversely, in the vCA1-spd adjacent to the SO, the expression levels of *Npy2r* were significantly higher than those of *Npy1r* (Fig. 4c, d and Extended Data Fig. 14g). Simultaneously examining the spatial expression pattern of *Npy* in the vCA1 using dFISH, we observed predominant arrangement of *Npy* in the SO proximate to vCA1-spd (Fig. 4c, e and Extended Data Fig. 14h).

**Fig 4.**
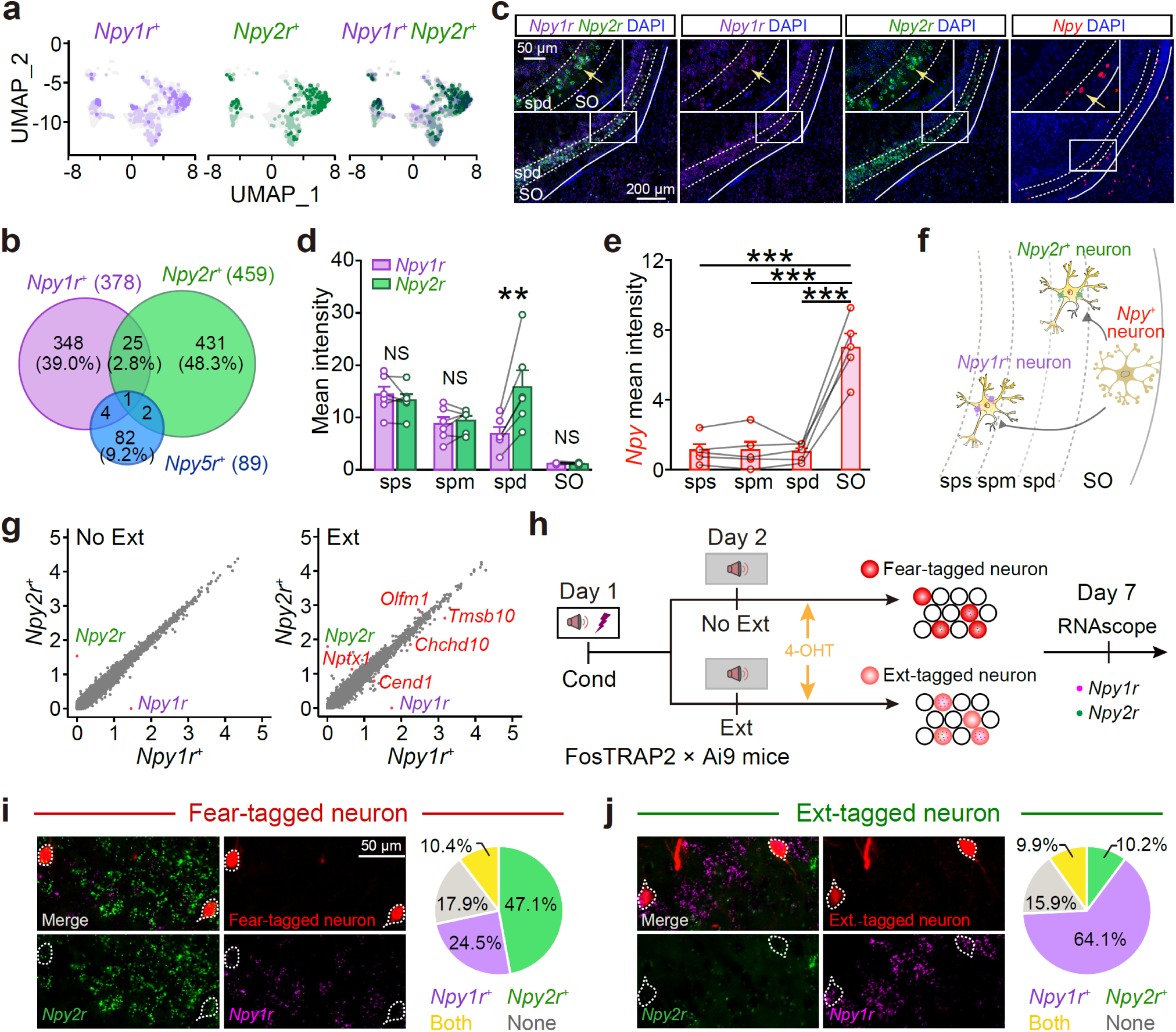
NPY bifurcates its roles in largely non-overlapping NPY receptor subtype-expressing neurons in the vCA1. **a**, UMAP resolved separated *Npy1r*-expressing and *Npy2r*-expressing clusters of vHPC excitatory and inhibitory neurons. **b**, Venn diagrams displayed the percentage distributions of *Npy1r*-, *Npy2r*-, and *Npy5r*-expressing neurons within NPY receptor-positive neurons in vHPC. The numbers indicate the percentage of *Npy1r*-, *Npy2r*-, and *Npy5r*-expressing neurons. **c**, Representative images showcasing *Npy1r*, *Npy2r*, and *Npy* distribution in vCA1. **d**, From the superficial (sps) to the middle (spm) to the deep (spd) vCA1 pyramidal layers, the expression of *Npy1r* gradually decreased; in the vCA1-spd adjacent to the SO, the expression levels of *Npy2r* were significantly higher than those of *Npy1r*. n = 6 mice. sps, NS, *P =* 0.298; spm, NS, *P* = 0.449; spd, \*\**P* = 0.007; SO, NS, *P* = 0.419, two-tailed paired Student’s *t*-test. **e**, Predominant arrangement of *Npy* in the SO proximate to vCA1-spd. n = 5 mice. SO *vs.* sps, \*\*\**P* = 0.0003; SO *vs.* spm, \*\*\**P* = 0.0003; SO *vs.* spd, \*\*\**P* = 0.0008, two-tailed paired Student’s *t*-test. **f**, A schematic diagram illustrating that NPY^+^ neurons bifurcate into distinct neurons expressing different NPY receptors in the vHPC, with closer proximity to the *Npy2r*-expressing neurons. **g**, During extinction, NPY released in the vCA1 primes differential transcriptional responses in neurons expressing *Npy1r* and *Npy2r*. **h**, Experimental procedure for labeling Fear-tagged and Ext-tagged neurons and determining the expression of *Npy1r* and *Npy2r* on these corresponding neurons. **i, j**, Quantification and comparison the distribution of *Npy1r* and *Npy2r* in Fear-(**i**) and Ext-tagged neurons (**j**). n = 4 and 3 mice for No Ext and Ext, respectively.

Notably, the NPY^+^ interneurons were physically closer to *Npy2r*-expressing neurons and farther away from *Npy1r*-expressing neurons in the vCA1 (Fig. 4c–f). Because of much higher affinity of NPY2r for NPY than NPY1r^43,44^, we postulated that the *Npy2r*-expressing neurons are preferentially inhibited as a result of the couplings of the NPY2r to Gi/o signaling pathway. Examining the single-cell transcriptome data of *Npy1r*- and *Npy2r*-expressing neurons to retrieve differentially expressed genes (DEGs), we identified two DEGs enriched in *Npy2r*-expressing neurons and three DEGs enriched in *Npy1r*-expressing neurons specifically under Ext but not No Ext conditions (Fig. 4g).

This signifies that NPY released during extinction in the vCA1 primes differential transcriptional responses in neurons expressing *Npy1r* and *Npy2r*, respectively, which likely represents two sub-ensembles of the same engram.

To further investigate the potential roles of putative *Npy1r*- and *Npy2r* sub-ensembles in fear- and/or extinction-engram neurons in the vCA1, we utilized the targeted recombination in active populations (TRAP) system^33,45^ with TRAP2::Ai9 (lox-stop-lox-tdTomato) double transgenic mice to tag active neurons during specific behavioral windows linked to fear or extinction, respectively. Fear memory retrieval (No Ext) was induced by exposing mice to 2 × CS with a high fear level, and intraperitoneal injections of 4-OHT to tag fear engrams (Fear-tagged neurons) with tdTomato fluorescence. Conversely, extinction learning (Ext) involved exposing mice to 20 × CS with a gradually decreasing fear level, and intraperitoneal injections of 4-OHT were given to tag extinction engrams (Ext-tagged neurons). *In situ* hybridization analysis (Fig. 4h) of *Npy1r* and *Npy2r* in either fear- or Ext-tagged neurons revealed that the percentage of *Npy2r*-expressing neurons was significantly higher than that of *Npy1r*-expressing neurons in fear-tagged neurons, while the order was opposite in Ext-tagged neurons (Fig. 4i, j). This raises an interesting possibility that NPY bifurcates its roles in differentially tagging two sub-ensembles of engram neurons for fear memory and extinction.

### Genetic deletion or pharmacological perturbations to *Npy1r* or *Npy2r* sub-ensembles selectively impact memory liability and stability

To explore the potential roles of *Npy1r* and *Npy2r* sub-ensembles, we utilized TRAP2::Cas9 (lox-stop-lox-Cas9) double transgenic mice in which the Cas9 protein was specifically expressed in Fear- or Ext-tagged neurons, inducing a double-strand break in the *Npy2r* or *Npy1r* genes targeted by corresponding guide RNAs. This process led to the formation of loss-of-function indel mutations by non-homologous end joining, enabling site-specific targeted mutation of the *Npy1r* or *Npy2r* genes *in vivo*, as confirmed by post hoc analyses (Extended Data Fig. 15). We assessed fear- and extinction-tagged ensembles after selectively knocking out *Npy1r* or *Npy2r* (Fig. 5a). For fear-tagged neurons, knocking out *Npy1r* impacted neither Stage 1 nor Stage 2 extinction, whereas knocking out *Npy2r* significantly slowed the rate of Stage1 and attenuated Stage 2 (Fig. 5b, c and Extended Data Fig. 16a–c), mirroring the effect of optogenetic inhibition of NPY^+^ interneurons (Fig. 3c, d). Leveraging the phenomenon of spontaneous recovery of extinguished fear over time^23,25^, we evaluated extinction relearning and retrieval after knocking out *Npy1r* or *Npy2r* in extinction-tagged neurons active during the first-round extinction learning. Knocking out *Npy1r* augmented both the rate and degree of extinction of both stages where on the contrary deletion of *Npy2r* had little effects on either stage (Fig. 5d, e and Extended Data Fig. 16d–f). These results suggested that NPY2R and NPY1R, within fear- and extinction-tagged ensembles, respectively, exhibit distinct effects on fear extinction, with one exerting a negative influence and the other a positive one.

**Fig 5.**
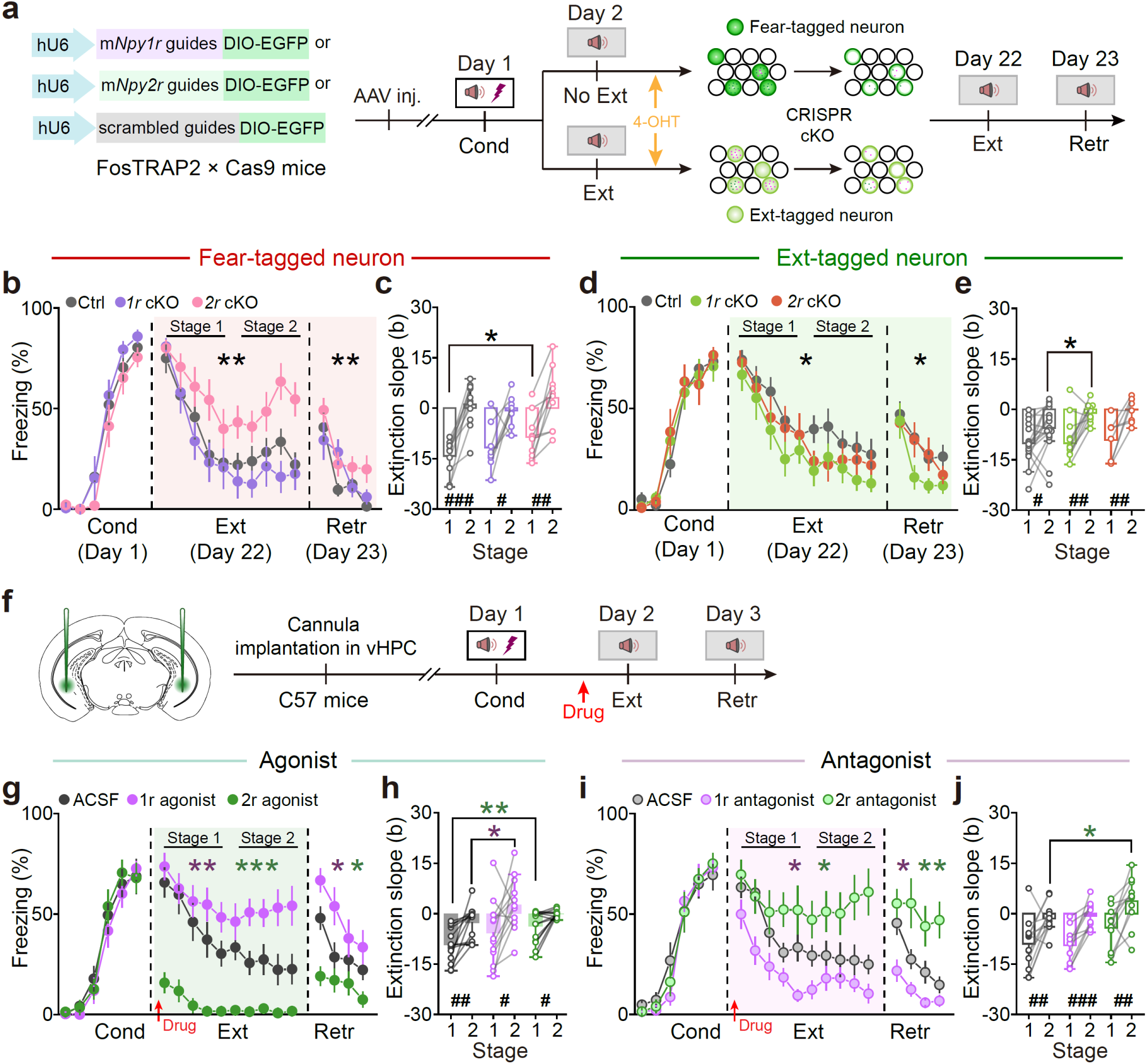
Genetic deletion or pharmacological perturbations to *Npy1r* or *Npy2r* sub-ensembles selectively impact memory liability and stability. a, Behavioral procedure, and schematic of *Npy2r*, *Npy1r* gene cKO in Fear- and Ext-tagged neurons respectively. **b, c**, *Npy2r* cKO in Fear-tagged neurons significantly slowed the rate of Stage1 and attenuated Stage 2, but not *Npy1r* cKO. n = 11 mice for Ctrl, 8 mice for *Npy1r* cKO, 10 mice for *Npy2r* cKO. (**b**) Two-way repeated ANOVA, main effects of group. Ext: F_(2, 26)_ = 4.973, \**P* = 0.015; Ctrl *vs. Npy1r* cKO, NS, *P* = 0.762; Ctrl *vs. Npy2r* cKO, \*\**P* = 0.007; *Npy1r* cKO *vs. Npy2r* cKO, \**P* = 0.023. Retr: F_(2, 26)_ = 4.170, \**P* = 0.027; Ctrl *vs. Npy1r* cKO, NS, *P* = 0.415; Ctrl *vs. Npy2r* cKO, \*\**P* = 0.009; *Npy1r* cKO *vs. Npy2r* cKO, NS, *P* = 0.082. (**c**) Stage 1 *vs.* Stage 2: Ctrl, ^###^*P* < 0.0001; *Npy1r* cKO, ^#^*P* = 0.015; *Npy2r* cKO, ^##^*P* = 0.002, two-tailed paired Student’s *t*-test. Ctrl *vs. Npy2r* cKO, \**P* = 0.041, unpaired Student’s *t*-test. **d, e**, *Npy1r* cKO in Ext-tagged neurons augmented both the rate and degree of extinction of both stages, but not *Npy2r* cKO. n = 17 mice for Ctrl, 12 mice for *Npy1r* cKO, 8 mice for *Npy2r* cKO. (**d**) Two-way repeated ANOVA, main effects of group. Ext: F_(2, 34)_ = 2.351, NS, *P* = 0.111; Ctrl *vs. Npy2r* cKO, NS, *P* = 0.316; Ctrl *vs.1r* cKO, \**P* = 0.038; *Npy1r* cKO *vs. Npy2r* cKO, NS, *P* = 0.417. Retr: F_(2, 34)_ = 2.833, NS, *P* = 0.073; Ctrl *vs. Npy2r* cKO, NS, *P* = 0.660; Ctrl *vs. Npy1r* cKO, \**P* = 0.025; *Npy1r* cKO *vs. Npy2r* cKO, NS, *P* = 0.139. (**e**) Stage 1 *vs.* Stage 2: Ctrl, ^#^*P* = 0.018; *Npy1r* cKO, ^##^*P* = 0.002; *Npy2r* cKO, ^##^*P* = 0.009, two-tailed paired Student’s *t*-test. Ctrl *vs. Npy1r* cKO, \**P* = 0.044, unpaired Student’s *t*-test. **f**, Schematic of cannula implantation and behavioral procedure with drugs microinjection into vHPC. **g, h**, The agonist for NPY2r caused excessive Stage 1 extinction with diminished Stage 2, while the agonist of NPY1r slightly slowed Stage 1 extinction and stabilized freezing at a much higher level than the control. n = 13 mice for ACSF, n = 13 mice for NPY1r agonist, n = 14 mice for NPY2r agonist. (**g**) Two-way repeated ANOVA. Ext: F_(2, 37)_ = 33.430, \*\*\**P* < 0.001; ACSF *vs.* NPY1r agonist, \*\**P* = 0.008; ACSF *vs.* NPY2r agonist, \*\*\**P* < 0.0001. Retr: F_(2, 37)_ = 7.411, \*\**P* = 0.002; ACSF *vs.* NPY1r agonist, \**P* = 0.040; ACSF *vs.* NPY2r agonist, \**P* = 0.035. (**h**) Stage 1 *vs.* Stage 2: ACSF, ^##^*P* = 0.003; NPY1r agonist, ^#^*P* = 0.023; NPY2r agonist, ^#^*P* = 0.011, two-tailed paired Student’s *t*-test. ACSF *vs.* NPY1r agonist, \**P* = 0.040; ACSF *vs.* NPY2r agonist, \*\**P* = 0.009, unpaired Student’s *t*-test. **i, j**, The antagonist of NPY1r promoted extinction, whereas that of NPY2r impeded it. n = 13 mice for ACSF, n = 10 mice for NPY1r antagonist, n = 11 mice for NPY2r antagonist. (**i**) Two-way repeated ANOVA. Ext: F _(2, 31)_ = 10.728, \*\*\**P* = 0.0003; ACSF *vs.* NPY1r antagonist, \**P* = 0.034; ACSF *vs.* NPY2r antagonist, \**P* = 0.013. Retr: F_(2, 31)_ = 13.544, \*\*\**P* < 0.0001; ACSF *vs.* NPY1r antagonist, \**P* = 0.041; ACSF *vs.* NPY2r antagonist, \*\**P* = 0.002. (**j**) Stage 1 *vs.* Stage 2: Ctrl, ^##^*P* = 0.006; NPY1r antagonist, ^###^*P* = 0.0005; NPY2r antagonist, ^##^*P* = 0.006, two-tailed paired Student’s *t*-test. ACSF *vs.* NPY2r antagonist, \**P* = 0.037, unpaired Student’s *t*-test.

To directly decipher the roles of *Npy1r* or *Npy2r* sub-ensembles in memory extinction, we locally delivered specific agonists and antagonists for NPY1r or NPY2r just before extinction learning through cannula implantation in the vCA1 (Fig. 5f). Notably, the agonist for NPY2r caused excessive Stage 1 extinction with diminished Stage 2, while the agonist of NPY1r slightly slowed Stage 1 extinction and stabilized freezing at a much higher level than the control, suggesting the role of NPY1r sub-ensemble being contingent upon that of NPY2r sub-ensemble (Fig. 5g, h and Extended Data Fig. 17a–c). Conversely, the antagonist of NPY1r promoted extinction, whereas that of NPY2r impeded it (Fig. 5i, j and Extended Data Fig. 17d–f), in line with the behavioral results from genetic deletion of *Npy2r* and *Npy1r* from neurons in two sub-ensembles as well as electrophysiology analyses of neuronal excitability and synaptic transmission in these mice (Extended Data Figs. 18 and 19). Pharmacological interventions targeting NPY5r in vCA1 had no discernible effects on fear extinction (Extended Data Fig. 20). These findings highlight distinctive actions of NPY1r and NPY2r in the fear extinction process and support the idea that NPY interneuron co-opts its roles in fear memory and extinction by activity-dependent release of NPY itself to sequentially act upon *Npy2r-* and *Npy1r*-expressing neurons, thereby permitting the transition from fear to extinction sub-ensembles. Specifically, NPY2r sub-ensemble engages in Stage 1 extinction, gating memory lability, while NPY1r functions in Stage 2 to determine the setpoint of memory stability during extinction (Extended Data Fig. 21).

## Discussion

The organization of engrams, specific neuronal ensembles recruited in an activity-dependent manner, is a fundamental process in memory storage and retrieval. IEGs, such as c-Fos and Arc, have been widely acknowledged as generic markers for activated neurons during learning, later identified as memory engrams^1,46^. Leveraging single-cell transcriptomic profiling, our study reveals a previously underappreciated population of vCA1 NPY^+^ neurons that exhibit exceptional peptide specificity to target specific sub-ensembles and expediate extinction learning by peptidergic transmission. The proportion and activity of NPY^+^ interneurons incrementally emerge among heterogeneous GABAergic inhibitory cell types in vCA1, locally driving NPY release onto *Npy1r*- and *Npy2r*-expressing neurons to regulate interconvertibility of memory engrams, underscoring the pivot roles of peptidergic GABAergic interneurons in determining engram lability and stability of memory.

Using genetic encoded sensors for NPY and calcium complemented by optogenetic and pharmacological manipulations, we demonstrated the neuron-type and region-specific regulation of NPY dynamics during extinction learning, as evidenced by tight correlation between NPY dynamics and the transition in behavioral states from fear-on to fear-off. Remarkably, optogenetic activation of vCA1 NPY^+^ interneurons, even in the absence of additional extinction learning, was proved sufficient to recapitulate the efficacy of behavioral extinction training, implicating the potential of vCA1 NPY^+^ interneurons as a cell-type-specific engram for fear extinction. It is noteworthy that NPY^+^ neurons located in the hypothalamic ARC region, another abundant source of central NPY^+^ neurons, did not exhibit extinction-dependent dynamics in neuronal activity or NPY release and fear extinction, reinforcing the notion that local release of NPY from vCA1 NPY^+^ neurons, but not global volume dispersion of NPY from ARC, gives rise to its specificity to selectively act on extinction memory engram.

Various GABAergic interneurons have been implicated in exerting direct inhibitory control over the original fear memory engrams^12,47,48^. The extinction-tagged neurons (extinction engrams), primarily composed of glutamatergic excitatory neurons^13^, directly excite the GABAergic inhibitory neurons to exert inhibitory control over the original fear memory engrams. However, these unidirectional control mechanisms may not fully explain the dynamic interactions between contrasting and interconvertible fear and extinction memories^19,20,22,23^. This study introduces a distinct engram construct for malleable fear and extinction memories, in which single peptide bifurcates distinct neuronal sub-ensembles. This engram organization features the NPY^+^ neurons together with the neighboring *Npy1r*- and *Npy2r*-expressing neurons in the vCA1. Although the vCA1 NPY^+^ interneurons are, in fact, also GABAergic, their peptidergic transmission in fear extinction is evidently crucial. A particularly intriguing aspect of our findings is the physical distance-dependent configuration observed between NPY^+^ neurons and two largely nonoverlapping NPY receptor subtype-expressing neurons for fear or extinction engrams. The NPY^+^ neurons are distributed in the SO close to the vCA1 where NPY receptor-expressing neurons exhibit specific cytoarchitectural distributions. *Npy1r*-expressing neurons are enriched in the superficial and middle pyramidal layers, while *Npy2r*-expressing neurons are located in the deep pyramidal layers closer to the SO. NPY2r also demonstrates a much higher affinity^43,44^ to NPY compared to NPY1r, suggesting that proximal NPY2r sub-ensemble is preferentially targeted during the early stage of extinction and inhibited by locally released NPY to support the inactivation of fear engram and the formation of extinction engram. As NPY ramps up during extinction learning, distal NPY1r sub-ensemble is recruited to prevent over-formation of the extinction engram during the late stage of extinction. Thus, the interplay between receptor affinity and physical distance emerges as critical factors for locally released NPY to determine the lability and stability of linked memory engrams in the vCA1.

Our study challenges the notion that engrams associated with specific memories are composed of active neurons at random. Specifically, we discover that *Npy2r*-expressing neurons are more likely associated with fear engrams, while *Npy1r*-expressing neurons are more likely to represent extinction engrams. These associations are ultimately shaped by the spatiotemporal dynamics of NPY surrounding them during memory and extinction learning. Our working model argues for a substantial revision of the merely activity-dependent memory engram allocation mechanism^49^. We propose that peptide tagging of engram cells by interneurons, as exemplified by NPY that sequentially activate NPY2r and NPY1r, to act on two sets of neurons in non-overlapping manner, are set by design . The sub-ensembles are spatiotemporally positioned to co-opt extinction process against the original fear memory, but does not permanently erase it^19,20,22,23^. This ideally annotates the dichotomous nature of engram lability and stability for memory.

In conclusion, our study highlights a nuanced interplay between neuropeptide signaling and neuronal activity dynamics, shaping the formation and maintenance of memory and extinction engrams in the vCA1. Specifically, the intricacies of NPY-mediated signaling in fear extinction holds the promise for the development of innovative treatments for anxiety-related disorders such as PTSD in the future.

## Methods

### Mice

All animal procedures were ethically approved by the Animal Ethics Committee of Shanghai Jiao Tong University School of Medicine and the Institutional Animal Care and Use Committee (Department of Laboratory Animal Science, Shanghai Jiao Tong University School of Medicine; Policy Number DLAS-MP-ANIM. 01–05). Mice were housed in groups under a 12-h light/dark cycle with unrestricted access to food and water. Adult male mice (7–12 weeks old, C57BL/6J background) were selected for all experiments. The study utilized the following mouse strains: NPY-Cre mice (stock no. 027851), Fos2A-iCreER (TRAP2) mice (stock no. 030323), lox-stop-lox-tdTomato (Ai9) reporter mice (stock no. 007909), NPY-hrGFP mice (stock no. 006417), and CRISPR/Cas9 knock-in mice (stock no. 024857) were obtained from the Jackson Laboratory. To obtain the *Npy* knockout mice (*Npy^−/−^*), littermate *Npy* knockdown mice (*Npy^+/−^*) and littermate control mice (*Npy^+/+^*) mice, we crossed the female *Npy^+/−^*mice with the male *Npy^+/−^* mice. *Npy^+/−^* mice were obtained from GemPharmatech Co., Ltd. (Nanjing, China). C57BL/6J mice were procured from the Shanghai Laboratory Animal Center (SLAC) at the Chinese Academy of Sciences (Shanghai, China).

### Fear conditioning, extinction, and extinction retrieval

All auditory fear conditioning and extinction procedures were conducted using the Ugo Basile Fear Conditioning System (UGO Basile S.R.L., Italy), following established protocols^24^. Initially, mice were acclimated and habituated to the conditioning chamber for three consecutive days. The conditioning chambers (17 × 17 × 25 cm), equipped with stainless-steel shocking grids, were linked to a precision feedback current-regulated shocker (UGO Basile S.R.L., Italy). During habituation and fear conditioning, the chamber walls were adorned with black-and-white checkered wallpapers (context A) and cleaned with 75% ethanol.

On Day 1, individual mice underwent conditioning in context A with five pure tones (Conditioned Stimulus, CS; 4 kHz, 76 dB, 20 s each) at fixed intervals (140 s), each coinciding with a foot shock (Unconditioned Stimulus, US; 0.5 mA, 2 s). Auditory tones and foot shocks were autonomously controlled by the ANY-maze software (version 7.20, Stoelting Co., USA). Following conditioning, mice were returned to their home cages 60 s after the final tone, with individual cage floors and walls sanitized with 75% ethanol.

Twenty-four hours post-conditioning, the no-extinction training group (No Ext) received 2-trial CS presentations, while the extinction training group (Ext) experienced 20-trial CS presentations during the same timeframe, without foot shock. Both No Ext and Ext tests occurred in a chamber with a gray, non-shocking plexiglass floor and dark gray wallpaper (context B). The testing environment underwent cleaning with a 4% acetic acid solution between individual mouse tests. To minimize CS anticipation during extinction, a distinct tone duration of 30 s (deviating from the 20-s conditioning duration) was used. Extinction training involved exposing mice initially trained in context A to 20 CS presentations (4 kHz, 76 dB, 30 s each) without foot shocks, conducted in context B on Day 2. For extinction retrieval on Day 3, mice underwent 8-trial CS presentations in context B. Both No Ext and Ext conditions, as well as extinction retrieval tests, combined 2-trial CS presentations into a single block.

Throughout testing, the chamber was placed in a sound-attenuating enclosure with a ventilation fan and a single house light (UGO Basile S.R.L., Italy). Mouse locomotion within the chamber was recorded using a near-infrared camera, analyzed in real-time by the ANY-maze software. The freezing score, a unit-less metric generated by the software, indicated freezing periods based on predefined threshold settings. A fear response was defined as measurable behavioral freezing, marked by a cessation of movement lasting more than 2 s. For animals with an optical fiber attached to the head, freezing during tone presentations was independently scored by an experimenter in a double-blind manner to mitigate potential interference from light stimuli during test sessions. The duration of freezing during each tone (CS) presentation was quantified for analysis.

### Generation of Single-Cell Suspensions

Two hours after undergoing fear memory retrieval (No Ext) or extinction learning (Ext), the mice were deeply anesthetized with 1% sodium pentobarbital and then decapitated. Brains were quickly dissected and chilled in ice-cold N-methyl-D-glucamine-artificial cerebrospinal fluid (NMDG-ACSF) containing the following components (in mM): 93 N-methyl-D-glucamine, 2.5 KCl, 1.2 NaH_2_PO_4_, 30 NaHCO_3_, 20 HEPES, 25 D-glucose, 5 Sodium ascorbate, 2 Thiourea, 3 Sodium pyruvate, 10 MgSO_4_, 11.9 N-Acetyl-L-cysteine, 1 CaCl_2_, and 6 ml HCl (pH 7.35–7.45). Coronal brain slices (400 μm thick) containing regions of the vHPC were cut using a vibratome (Leica VT1000S, Germany), then promptly transferred to ice-cold NMDG-ACSF and incubated for 15 minutes. Actinomycin D (8 μM; Sigma-Aldrich, Catalog no. A1410) and Triptolide (10 μM) were added in this step to minimize artificially induced activation of early immediate genes. The region of vHPC were carefully microdissected using fine tweezers on ice. After tissue collection, it was transferred to an Eppendorf tube containing a digestion solution. This solution consisted of ACSF supplemented with 1 mg/ml pronase (Sigma, Catalog no. P6911-1G), ActD (8 μM), Triptolide (10 μM), TTX (1 μM) and D-APV (100 μM), and the tissue was digested for 20 min at 34 °C. The tissue was pipetted periodically every 10 min during this digestion. Then, the cell suspension was transferred to Sterile Earle’s Balanced Salt (EBSS) Solution (JinpanWorthington, Catalog no. LK003150) supplemented with ActD (8 μM), Triptolide (10 μM), TTX (1 μM) and D-APV (100 μM). Finally, it was washed with washing solution consisting of ACSF supplemented with ActD (8 μM), Triptolide (10 μM), TTX (1 μM) and D-APV (100 μM). All solutions were continually bubbled with O_2_ and CO_2_ (95% O_2_/5% CO_2_, v/v).

### Single-cell library preparation and sequencing

Single cell RNA-seq library were prepared from the single cell suspensions via the chromium single cell gene expression system (Chromium Single Cell 3’ Reagent Kits v2, 10x genomics), using the default protocols provided by 10 × genomics. The NovaSeq platforms were used for sequencing libraries.

### Genome alignment and Seurat analysis

Fastq files were aligned to the mouse reference genome mm10 and converted into gene expression matrices using the 10x Genomics Cell Ranger software (v3.0.0). Cells with less than 700 transcripts or 200 genes or more than 15000 transcripts or 5% of mitochondrial expression were filtered out as low-quality cells. To visualize the data, dimension reduction and clustering analysis were performed on the raw count matrices using the Seurat (v4.3.0) package in R (v4.1.3)^50^. The FindIntegrationAnchors and IntegrateData functions were used to integrate the No Ext and Ext datasets. The data were reduced using principal component analysis (PCA). The clusters were obtained using the FindNeighbors and FindClusters functions with the resolution set to 0.5, resulting in 35 cell clusters. The cluster marker genes were found using the FindAllMarkers function. The cell types were annotated by overlapping the cluster markers with the known cell-type marker genes.

Uniform Manifold Approximation and Projection (UMAP) dimensional reduction was further employed to visualize the cell classifications, resulting in 9 cell types. Differential expression analysis between the No Ext and Ext was carried out using the FindMarkers function with the method of MAST^51^. When identifying DEGs within different cell types and different clusters, we used a threshold of |log_2_FC| > 0.585 and *P* < 0.05.

### Virus constructs

The following viruses were utilized: AAV-EF1α-DIO-NpHR3.0-EGFP (Serotype 2/9), AAV-EF1α-DIO-ChR2-EGFP (Serotype 2/9), AAV-EF1α-DIO-EGFP (Serotype 2/9) were procured from Obio Technology Co., Ltd. (Shanghai). AAV-EF1α-DIO-ChR2-E123T/T159C-mCherry (Serotype 2/9), AAV-EF1α-DIO-mCherry (Serotype 2/9), AAV-hSyn-DIO-ChrimsonR-mCherry (Serotype 2/9), and AAV-hSyn-DIO-hM4Di-mCherry (Serotype 2/9) were obtained from Brain VTA (Wuhan, China). AAV-hSyn-GRAB_NPY1.0_-EGFP (Serotype 2/9), AAV-hSyn-GRAB_NPYmut_-EGFP (Serotype 2/9), and AAV-hSyn-GRAB_GABA0.8_-EGFP (Serotype 2/9) were purchased from WZ Biosciences Inc (Jinan, China). AAV2/9-3x(hU6-sgRNA.sp)(m*Npy1r*)-hEF1a-DIO-EGFP, AAV2/9-3x(hU6-sgRNA.sp)(m*Npy2r*)-hEF1a-DIO-EGFP, AAV2/9-hU6-sgRNA-hEF1a-DIO-EGFP, AAV2/9-H1-sgRNA.sp(m*slc32a1*)x3-CAG-DIO-mCherry, AAV2/9-H1-sgRNA.sp(m*Npy*)x3-CAG-DIO-mCherry, and AAV2/9-H1-sgRNA.sp(NC)-CAG-DIO-mCherry were generated by Shanghai Taitool Bioscience Co., Ltd. (Shanghai). All viral vectors were stored in aliquots at –80°C until further use, with viral titers for injection exceeding 10^12^ viral particles per ml.

### Surgery

Mice, aged 7–8 weeks, were anesthetized with a 1% sodium pentobarbital solution administered via a single intraperitoneal injection (10 ml per kg of body weight). After anesthesia, each mouse was immobilized in a stereotactic frame using non-rupture ear bars (RWD Life Science, Shenzhen, China). A midline scalp incision was made, and small bilateral craniotomies were created using a microdrill with 0.5-mm burrs. Glass pipettes (tip diameter: 10–20 μm) were crafted using a P-1000 Micropipette Puller (Sutter glass pipettes, Sutter Instrument Company, USA) for AAV microinjections. Initially filled with silicone oil, the microinjection pipettes were connected to a microinjector pump (RWD Life Science, Shenzhen, China) to ensure complete air exclusion. AAV-containing solutions were loaded into the pipette tips and injected at specified coordinates (in mm): vCA1 (anteroposterior to bregma, AP, −3.14; lateral to the midline, ML, ±3.20; below the bregma, DV, −4.00) and ARC (AP, −1.70; ML, ±0.30; DV, −5.80). Virus-containing solutions were injected bilaterally into the vCA1 (0.4 μl/side) and unilaterally into the ARC (0.3 μl/side) at a rate of 0.1 μl/min. After injection, the pipette remained in place for an additional 10 minutes to allow for adequate diffusion of the injectant. Mice were given a minimum of two weeks to recover before undergoing behavioral and other tests, and injection sites were examined at the experiment’s conclusion by assessing the expression of fluorescent proteins, such as EGFP or mCherry.

In optogenetic experiments, ceramic fiber optic cannulas (200 μm in diameter, 0.37 numerical aperture (NA), Hangzhou Newdoon Technology) were surgically implanted above the vCA1 (coordinates: AP, –3.14; ML, ±3.20; DV, –3.90) and ARC (coordinates: AP, –1.70; ML, ±0.30; DV, –5.70). These cannulas were secured in place using acrylic dental cement and skull screws.

### Optogenetic manipulations

A wired optogenetic system was utilized to modulate neuronal activity during behavioral assays, employing a light-emitting diode (LED) (Hangzhou Newdoon Technology Co. Ltd) connected to the optic patch cord through connectors on each end. Blue light (470 nm, 4–5mW) was delivered in 10-ms pulses at 20 Hz during extinction training. For optogenetic inhibition, red light (638 nm, 8–10 mW) was consistently administered. The duration of light pulses exceeded 5 s both before and after the CS presentations to ensure complete CS exposure during extinction training (30-s duration for each tone). To activate NPY^+^ neurons during freezing in response to CS presentations during extinction, if a fear response was detected by the ANY-maze software, blue light (470 nm, 4–5mW) was automatically triggered, and the blue light switched off when the fear response ceased. The ultimate output power varied depending on the light transmission efficacy of the optical fiber used.

### Open field test

Mice were acclimated to the experimental room from their home cages and given a minimum of 1 h for habituation before the start of the experiment. Individual mice were then placed in the outer area of a square Plexiglas open-field apparatus (40 × 40 × 35 cm), divided into a central field (20 × 20 cm) and an outer field. The total distance traveled was quantified using Noldus EthoVision XT (version 16.0, Noldus Information Technology, Netherlands). NPY-Cre mice underwent injections of DIO-ChR2, -NpHR, or -Control virus to modulate the activity of NPY^+^ neurons, with customized light stimulation protocols for either activation or inhibition of these neurons. The open-field test consisted of an 18-min session divided into six 3-min epochs, alternating between light off and light om periods, starting with a light-off epoch. For analysis and charts representing only “off” and “on” conditions, the three “off” epochs and three “on” epochs were combined, respectively. To ensure consistency, the open-field arena was thoroughly cleaned with 70% ethanol between each set of tests.

### Fiber photometry

Fiber photometry experiments were performed using a system from Thinker Tech (Nanjing, China). Fluorescent signals, generated by a 488-nm laser (OBIS 488LS; Coherent), were reflected by a dichroic mirror (MD498; Thorlabs), focused by a 10× objective lens (NA = 0.3; Olympus), and coupled to a rotary joint (FRJ_1 × 1_FC-FC, Doric Lenses, Canada). An optical fiber (200 mm O.D., NA = 0.37) was implanted into the vCA1 or ARC during virus injection and remained in place for two weeks for virus expression. Laser power at the fiber tip was adjusted to 40–50 μW to minimize bleaching of GCaMP6m probes or GRAB_NPY1.0_/GRAB_NPYmut_ sensors. Excitation fluorescence was collected by the same multi-mode optical fiber and converted into electrical signals by low-light detectors at the detection end to capture neural activity information. Signals were digitized at 100 Hz using a Power 1401 digitizer and Spike 2 software (CED, Cambridge, UK). Throughout behavioral tests, including fear learning, extinction training, or extinction retrieval, fluorescent intensities of GCaMP6m or sensors were recorded, with the pre-sound signal serving as the baseline. Average Ca^2+^ or sensor responses were computed using MATLAB, and data were exported to MATLAB mat files from Spike2 for further analysis. Permutational tests were employed for statistical significance, and ΔF/F values were visualized as heatmaps or per-event plots, with shaded areas indicating the standard error of the mean (SEM). The averaged ΔF/F during CS presentation and freezing/no-freezing states was defined, and the area under the curve (AUC, ΔF/F × s) was calculated for the same dataset.

### Cannula implantation and local drug injection

Mice were anesthetized with 1% sodium pentobarbital and securely fixed on a stereotaxic apparatus (RWD Life Science, Shenzhen, China). Stainless steel guide cannulas (RWD Life Science, Shenzhen, China) were bilaterally implanted into the vCA1, with the cannula tips positioned at the following coordinates (in mm): AP, –3.14; ML, ±3.20; DV, –3.90. The cannulas were firmly attached to the skull using acrylic cement and two skull screws. Stainless steel obturators (33 gauges) were inserted into the guide cannulas to prevent obstruction until drug infusion. Animals were allowed a 2-week recovery period post-surgery before undergoing behavioral tests. Mice were familiarized with the infusion procedure three days prior to drug injection. During drug infusion, mice were briefly head-restrained, the stainless-steel obturators were removed, and injection cannulas (33 gauges, RWD Life Science, Shenzhen, China) were inserted into the guide cannulas.

The injection cannulas protruded 0.50 mm from the guide cannula tips. The infusion cannula was connected via PE20 tubing to a microsyringe driven by a microinfusion pump (KDS 310, KD Scientific, USA). A total of 0.5 μl of drugs were bilaterally infused into the vCA1 at a controlled flow rate of 0.1 μl per minute. After completing the drug injection, the injection cannulas were left in place for 2 min to facilitate solution diffusion from the cannula tips. Subsequently, the stainless-steel obturators were reinserted into the guide cannulas, and the mice were returned to their home cages for a 15-min recovery period before behavioral tests.

### Drugs and concentrations

Neuropeptide Y (Tocris, Catalog no. 1153). NPY1r agonist, [Leu31, Pro34]-Neuropeptide Y (porcine) (MedChemExpress, Catalog no. HY-P0208). NPY2r agonist, N-Acetyl-[Leu^28^, Leu^31^]-Neuropeptide Y Fragment 24-36 (Sigma, Catalog no. N9398). NPY5r agonist, [cPP1-7, NPY19-23, Ala31, Aib32, Gln34]-hPancreatic Polypeptide (MedChemExpress, Catalog no. HY-P1324). The peptide and peptide agonists for NPY receptors were applied at a concentration of 0.25 mM for 0.5 μl. NPY1r antagonist, BIBO 3304 trifluoroacetate (Tocris, Catalog no. 2412) was applied at a concentration of 0.4 mM for 0.5 μl. NPY2r antagonist, BIIE 0246 hydrochloride (Tocris, Catalog no. 7377) was applied at a concentration of 2 mM for 0.5 μl. NPY5r antagonist, CGP 71683 hydrochloride (Tocris, Catalog no. 2199) was applied at a concentration of 5 mM for 0.5 μl.

### Engram labeling

Activity-dependent recombination was triggered using 4-hydroxytamoxifen (4-OHT, Sigma-Aldrich, Catalog no. H6278, USA). A 20 mg/ml solution of 4-OHT was prepared in ethanol by shaking at 37°C for 30 min. Subsequently, twice the volume of corn oil (Sigma-Aldrich, Catalog no. C8267, USA) was added to achieve a final concentration of 10 mg/ml of 4-OHT, and the ethanol was removed by vacuum under centrifugation. All injections were administered intraperitoneally.

Mice were transferred from the vivarium to an adjacent holding room at least 3 h before the test. For fear and extinction memory, activity-dependent neuronal tagging was initiated by a single intraperitoneal injection of 4-OHT (20 mg/kg for mice) after exposure to 2-trail CS or 20-trail CS, respectively. Subsequently, mice were returned to the vivarium, maintaining a regular 12-h light/dark cycle for the remainder of the experiment.

### Slice electrophysiology

Whole-cell recordings were conducted in acute brain slices obtained from behaviorally trained mice. Mice were deeply anesthetized with 1% sodium pentobarbital and subsequently decapitated. Brains were quickly dissected and chilled in ice-cold N-methyl-D-glucamine-artificial cerebrospinal fluid (NMDG-ACSF) containing the following components (in mM): 93 N-methyl-D-glucamine, 2.5 KCl, 1.2 NaH_2_PO_4_, 30 NaHCO_3_, 20 HEPES, 25 D-glucose, 5 Sodium ascorbate, 2 Thiourea, 3 Sodium pyruvate, 10 MgSO_4_, 11.9 N-Acetyl-L-cysteine, 1 CaCl_2_, and 6 ml HCl (pH 7.35–7.45). Coronal brain slices (300 μm thick) containing regions of the vCA1 were cut with a vibratome (Leica VT1000S, Germany). Slices were initially recovered for 5–10 minutes in NMDG-ACSF at 31°C. They were then transferred to another ACSF solution (in mM): 125 NaCl, 2.5 KCl, 12.5 D-glucose, 1 MgCl_2_, 2 CaCl_2_, 1.25 NaH_2_PO_4_, and 25 NaHCO_3_ (pH 7.35–7.45) for an additional 2 hours of recovery at 31°C before recordings. The slice was subsequently transferred to a recording chamber and continuously superfused with ACSF at a rate of 1–2 ml per minute. All solutions were continually bubbled with O_2_ and CO_2_ (95% O_2_/5% CO_2_, v/v). Neurons in the vCA1 were patched under visual guidance using infrared differential-interference contrast microscopy (BX51WI, Olympus, Japan), and an optiMOS camera (QImaging, Teledyne Imaging Group, Canada). Whole-cell patch-clamp recordings were performed using an Axon 200B amplifier (Molecular Devices, USA). Membrane currents and potentials were sampled and analyzed using a Digidata 1440 interface and a personal computer running Clampex and Clampfit software (pCLAMP 10.5, Molecular Devices, USA). Access resistance was maintained between 10–20 MΩ, and only cells with a change in access resistance <20% were included in the analysis. In specific recording situations, NPY (1 μM) was added to the ACSF to activate NPY receptors in the vCA1.

#### Spike firing

The spiking activity and membranous properties of cell populations in the vCA1 were quantified using an internal solution comprising the following concentrations (in mM): 145 potassium gluconate, 5 NaCl, 10 HEPES, 2 MgATP, 0.1 Na_2_GTP, 0.2 EGTA, and 1 MgCl_2_ (280– 300 mOsm, pH 7.2 with KOH). Subsequent data analysis was performed using the MiniAnalysis Program (Version 6.0.1, Synaptosoft, USA) with an amplitude threshold set at 40 mV.

#### Spontaneous excitatory postsynaptic currents (sEPSCs)

For recording sEPSCs in cell populations within the vCA1, a holding potential of –70 mV was maintained. Patch pipettes were filled with a Cs^+^-based solution containing the following concentrations (in mM): 132.5 Cs-gluconate, 17.5 CsCl, 2 MgCl_2_, 0.5 EGTA, 10 HEPES, 2 Na_2_ATP, with pH adjusted to 7.3 using CsOH, and osmolarity set at 280–290 mOsm. sEPSCs were recorded for 5-10 min and analyzed from 300 to 400 s after the establishment and stabilization of the recording. Data analysis was conducted using the Mini-analysis Program (Synaptosoft) with an amplitude threshold set at 10 pA, while other parameters remained at their default values.

#### Optical stimulation response

Optical stimulation of ChR2- or NpHR-expressing neurons was conducted using a collimated LED (Lumen Dynamics Group Inc, USA) with peak wavelengths of 473 or 638 nm, respectively. The LED was connected to an Axon 200B amplifier to initiate photostimulation. The brain slice in the recording chamber was illuminated through a 40× water-immersion objective lens (LUMPLFLN 40XW, Olympus, Japan). The intensity of photostimulation was directly regulated by the stimulator (2–18 mW/mm^2^), while the duration was set through Digidata 1440 and pClamp 10.5 software. The functional potency of the ChR2-expressing virus was validated by measuring the numbers of action potentials (APs) elicited at different frequencies of blue-light stimulation (1 ms; 5, 10, and 20 Hz) and the inward photocurrents (500-ms pulse) mediated by ChR2 in brain slices. To validate the functional efficacy of NpHR-mediated optogenetic inhibition, red light (638 nm, 500-ms pulse) was administered to attenuate spikes under current clamp mode and induce the outward photocurrents (500-ms pulse) mediated by NpHR.

### CRISPR-mediated gene knockout

The CRISPR-associated endonuclease Cas9 from Streptococcus pyogenes (SpCas9) was employed to induce indels in the *Npy* (NC_109648), *slc32a1* (NC_22348), *Npy1r* (NC_18166), and *Npy2r* (NC_18167) genes. Corresponding single guide RNAs (sgRNAs) for each gene were designed and produced by Taitool Bioscience Co. Ltd. using the GRCM38 (Mus musculus) reference genome. Ten sgRNAs were selected for each gene based on computed specificity and efficiency scores (CHOPCHOP). Subsequently, these sgRNAs were cloned into a 2-in-1 reporter vector (CRPT001, Taitool), individually expressing a single sgRNA along with a cut-activated EGFP gene. The sgRNA target sequence was positioned ahead of the EGFP ORF within the vector. The reporter vectors were co-transfected with a Cas9-expressing plasmid into 293T cells. Functional Cas9/sgRNA complexes would cut the target on the reporter vector and rescue the expression of EGFP. Subsequently, the top three sgRNAs were selected and cloned into an adeno-associated virus (AAV) vector tandemly. The chosen sgRNAs were as follows:

5’-CGTATGCACCCACCTCTGTC-3’, 5’-AACAAGCGAATGGGGCTGTG-3’, and 5’-CAGAAAACGCCCCCAGAACA-3’ for the *Npy* gene targeting exon 2 of the reverse strand, exon 2 of the forward strand, and exon 3 of the forward strand of the *Npy* coding region, respectively.

5’-GCCACCGATGAGGAAGCGGT-3’, GCGCGTGCGGGACTCGTATG and 5’-GCACGCGATGAGGATCTTGC-3’ for *slc32a1* gene to target exon 2 of the forward strand, exon 3 of the forward strand and exon 3 of the reverse strand of the *slc32a1* coding region.

5’-GATGGAGACACACTGTACAA, GATGGACCACTGGGTCTTCG-3’, and 5’-GGCCGAAATACTGCAGCACC-3’ for the *Npy1r* gene targeting exon 2 of the reverse strand, exon 2 of the forward strand, and exon 2 of the reverse strand of the *Npy1r* coding region, respectively.

5’-GTTGGATGCCATTCACTCGG-3’, 5’-GAATCCAGGAATTTCAAGCA-3’, and 5’-GAGAAAGATATGATGCCCAG-3’ for the *Npy2r* gene targeting exon 2 of the forward strand, exon 1 of the reverse strand, and exon 2 of the reverse strand, respectively (according to transcript variant 1).

A non-targeting negative control guide was designed with a scrambled sequence with no known targets in the genome (5’-GCTGAGTACTTCGAAATGTC-3’). sgRNAs for *Npy* and *slc32a1* was cloned into the pAAV2-H1-sgRNA.sp(MCS)x3-CAG-DIO-mCherry-WPRE-pA vector (AC610-C, Taitool), while sgRNAs for *Npy1r* or *Npy2r* were cloned into the pAAV-(U6-sgRNA.sp(MCS)) x 3-hEF1a-DIO-EGFP-WPRE-pA vector (AC705-C, Taitool). These plasmids were co-transfected into HEK293 cells with AAV9 Cap. Rep and adenovirus helper to produce recombinant AAV vectors. Finally, AAV vectors were administered via stereotactic injection.

### In situ hybridization

Fluorescent in situ hybridization was conducted using the ACDBio V2 RNAScope kit (Advanced Cell Diagnostics). The following products were employed: RNAscope Multiplex Fluorescent Detection Reagent V2 (Catalog no. 323110), Pretreatment Reagents (Catalog no. 322381 and 322000), RNAscope Wash Buffer Reagents (Catalog no. 310091), probes for *Npy* (Catalog no. 313321-C1), *slc32a1* (Catalog no. 319191-C3), *slc17a7* (Catalog no. 416631-c2), c-fos (Catalog no. 316921-C2), *Npy1r* (Catalog no. 427021-C3), and *Npy2r* (Catalog no. 315951-C2), along with the PerkinElmer TSA Plus Fluorescence Palette Kit (Catalog no. NEL760001KT). Mice were deeply anesthetized with 1% sodium pentobarbital and transcardially perfused with ice-cold PBS before being fixed in 4% paraformaldehyde for 24 hours at 4 °C. After dehydration, brain tissue was embedded in optimal cutting temperature (OCT) and frozen at -80 °C. Coronal brain slices encompassing the entire vCA1 were sectioned at a thickness of 14 μm using a cryostat (Leica CM 1950). Slides were baked for 30 minutes at 60°C. Subsequently, 5–8 drops of Hydrogen Peroxide were added to cover the entire section, followed by an incubation for 10 minutes at room temperature (RT). The slides were then washed three times in distilled water. The tissues were brought to 1X Target Retrieval Reagent and maintained for 5 minutes at 99-100 °C. Following this, the slides were washed three times in distilled water. They were then transferred to 100% alcohol for 3 minutes.

Approximately 5 drops of Protease III were added to each section, and the samples were incubated for 30 minutes at 40°C. Excess liquid was removed from the slides, and 4–6 drops of the appropriate probe mix were added to entirely cover each slide at 40 °C for two hours. To amplify the signals, tissues were sequentially immersed in AMP1, AMP2, and AMP3 at 40 °C. AMP1 and AMP2 were incubated for 30 minutes each, while AMP3 was incubated for 15 minutes. After each incubation, the appropriate HRP channel was chosen, and 4–6 drops of HRP-C1 or HRP-C2 or HRP-C3 were added to entirely cover each slide at 40 °C for 15 minutes, followed by incubation with HRP blocker for 15 minutes at 40 °C. For a specific HRP channel, corresponding diluted Opal™ Dye was added to each slide and incubated for 30 minutes at 40°C. Npy was labeled in green (Opal™ 520) or red (Opal™ 570), *Npy1r* was labeled in far red (Opal™ 690), c-fos was labeled in red (Opal™ 570), and *Npy2r*, *slc32a1*, and *slc17a7* were labeled in green (Opal™ 520). Starting with protease III treatment, between all steps, slides were washed three times in 1× RNAscope wash buffer for 2 minutes.

Subsequently, the slides were incubated in ACDBio DAPI for 10 minutes, washed, dried for 20 minutes, cover-slipped with mounting media (SouthernBiotech, 0100-01), and left to dry overnight before imaging.

### Histology and fluorescent immunostaining

Fluorescent in situ hybridization was performed using the ACDBio V2 RNAscope kit (Advanced Cell Diagnostics, USA). The following products were utilized: RNAscope Multiplex Fluorescent Detection Reagent V2 (Catalog no. 323110), Pretreatment Reagents (Catalog no. 322381 and 322000), RNAscope Wash Buffer Reagents (Catalog no. 310091), probes for *Npy* (Catalog no. 313321-C1), *slc32a1* (Catalog no. 319191-C3), *slc17a7* (Catalog no. 416631-c2), *c-fos* (Catalog no. 316921-C2), *Npy1r* (Catalog no. 427021-C3), and *Npy2r* (Catalog no. 315951-C2), along with the PerkinElmer TSA Plus Fluorescence Palette Kit (Catalog no. NEL760001KT).

Mice were deeply anesthetized with 1% sodium pentobarbital and transcardially perfused with ice-cold PBS before fixation in 4% paraformaldehyde for 24 h at 4°C. Following dehydration, brain tissue was embedded in optimal cutting temperature (OCT) and frozen at –80°C. Coronal brain slices encompassing the entire vCA1 were sectioned at a thickness of 14 μm using a cryostat (Leica CM 1950, Japan). Slides were baked for 30 minutes at 60°C. Subsequently, 5–8 drops of Hydrogen Peroxide were added to cover each entire brain slice, followed by an incubation for 10 min at room temperature (RT). The slides were then washed three times in distilled water. The tissues were brought to 1X Target Retrieval Reagent and maintained for 5 minutes at 99–100°C. Following this, the slides were washed three times in distilled water. They were then transferred to 100% alcohol for 3 min. Approximately 5 drops of Protease III were added to each slice, and the samples were incubated for 30 min at 40°C. Excess liquid was removed from the slices, and 4–6 drops of the appropriate probe mix were added to entirely cover each slice at 40°C for 2 h. To amplify the signals, tissues were sequentially immersed in AMP1, AMP2, and AMP3 at 40°C. AMP1 and AMP2 were incubated for 30 min each, while AMP3 was incubated for 15 min. After each incubation, the appropriate HRP channel was chosen, and 4–6 drops of HRP-C1 or HRP-C2 or HRP-C3 were added to entirely cover each slice at 40°C for 15 min, followed by incubation with HRP blocker for 15 min at 40°C. For a specific HRP channel, corresponding diluted Opal^TM^ Dye was added to each slice and incubated for 30 minutes at 40°C. Npy was labeled in green (Opal^TM^ 520) or red (Opal^TM^ 570), Npy1r was labeled in far red (Opal^TM^ 690), c-fos was labeled in red (Opal^TM^ 70), and *Npy2r*, *slc32a1*, and *slc17a7* were labeled in green (Opal^TM^ 20). Starting with protease III treatment, between all steps, slides were washed three times in 1× RNAscope wash buffer for 2 min.

Subsequently, the slides were incubated in ACDBio DAPI for 10 min, washed, dried for 20 min, cover-slipped with mounting media (SouthernBiotech, 0100-01), and left to dry overnight before imaging.

### Statistical analyses

Statistical analyses were conducted using IBM SPSS Statistics 25, Origin 2022 Software, and Office 2019 (Microsoft, USA). The graphs were generated using Origin Software. Data are presented as the mean ± SEM unless indicated otherwise. Most histograms display individual data points representing the values and numbers of individual samples for each condition. Data distributions were tested for normality, and homogeneity of variance among groups was assessed using Levene’s test. Statistical comparisons were carried out using two-tailed Student’s *t*-test, as well as one-way analyses of variance (ANOVAs) or two-way repeated measures ANOVAs. For post hoc analysis, Bonferroni’s corrections for multiple comparisons were applied. A significance level of P < 0.05 was considered statistically significant. Significance is primarily denoted as \**P* < 0.05, \*\**P* < 0.01, and \*\*\**P* < 0.001. In some cases, it is indicated as ^#^*P* < 0.05, ^##^*P* < 0.01, and ^###^*P* < 0.001 for multiple comparisons. NS, denotes non-significant values.

## Data and code availability

All data needed to evaluate the conclusions in the paper are present in the paper and/or the Supplementary Materials. Additional data available from authors upon request.

## Acknowledgments

This study was supported by grants from the STI2030-Major Projects (2021ZD0202800), the National Natural Science Foundation of China (31930050, 81961128024, 32071023, 32200821, 32371078, and 32300843), the Science and Technology Commission of Shanghai Municipality (22XD1420700 and 23YF1433900), the Shanghai Municipal Health Commission (2022XD046), and innovative research team of high-level local universities in Shanghai. The *Npy^+/−^* mice were kindly shared by Dr. Xiajing Tong’s group at ShanghaiTech University.

## AUTHOR CONTRIBUTIONS

Y.-J.W., X.G., Q.L., W.-G.L., and T.-L.X. conceived the project, designed the experiments, and interpreted the results. Y.-J.W. and X.G. performed the majority of behavioral experiments, animal surgery, immunohistochemistry, and data analysis. M.X., Q.W., X.Y., Z.-J.L., Z.-H.J., H.C., X.-Y.Z., and X.B. assisted with some of the behavioral experiments and conducted viral injections. Q.J. and Ying Li performed genotyping. Y.-J.W., H.W., and Yulong Li performed NPY dynamics measurement. Y.-J.W., X.G., Y.K., S.Y., J.H., and Q.L. did sing-cell preparation and analyses. M-X.Z. and L.-Y.W. contributed to experimental design and data interpretation. Y.-J.W., L.-Y.W., W.-G.L. and T.-L.X. wrote the manuscript with contributions from all authors. All authors read and approved the final manuscript.

## DECLARATION OF INTERESTS

The authors declare no competing interests.

## Additional information

Extended data includes 21 Supplementary figures and their legends are available for this paper online.

## Extended Data

**Extended Data Fig 1.**
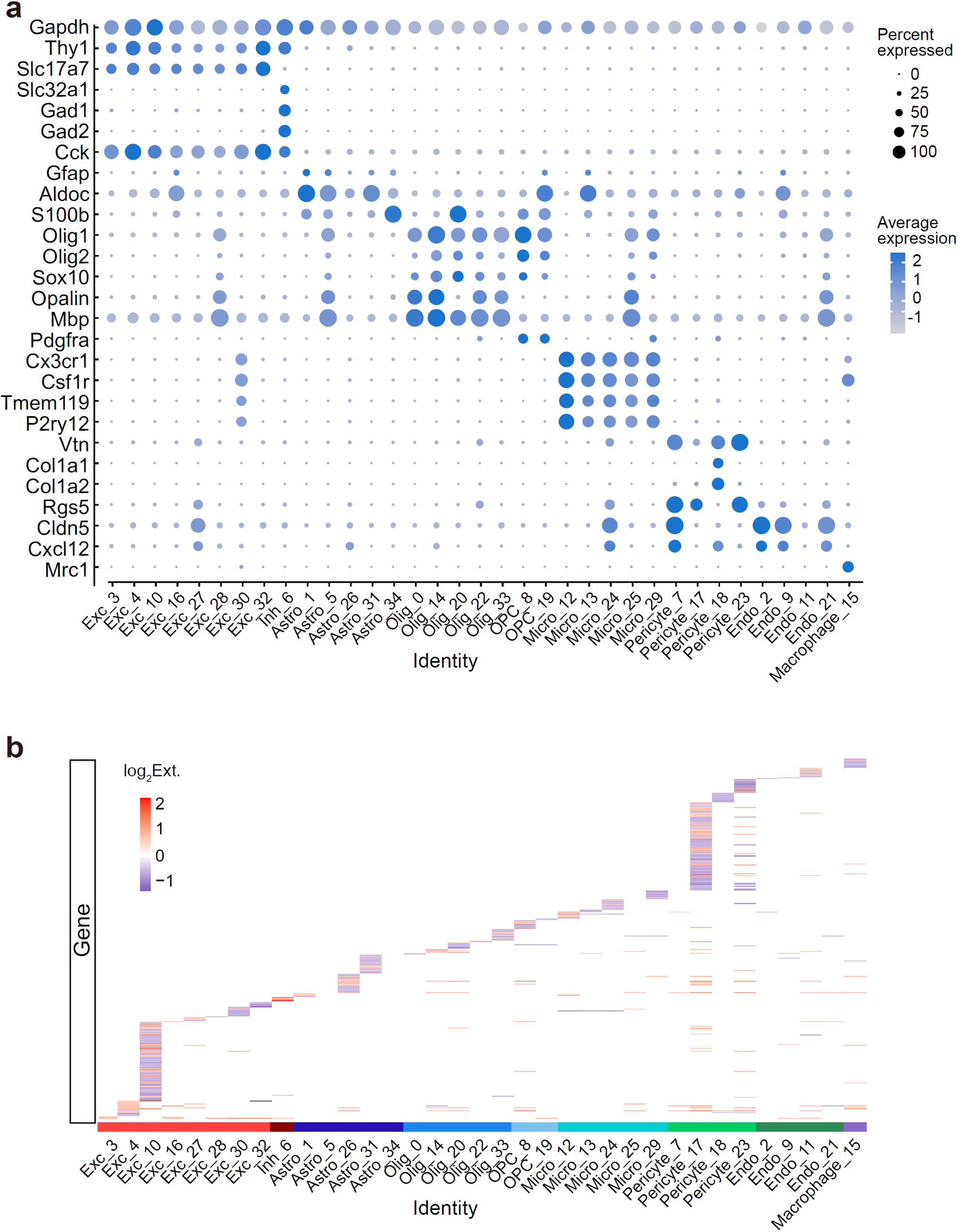
Molecular signatures and DEGs of different clusters in vHPC. **a**, Molecular signatures of different clusters by percentage of cells expressing the gene (circle size) and average gene expression level (color scale). These clusters encompassed excitatory neurons (Clusters 3, 4, 10, 16, 27, 28, 30, 32), inhibitory neurons (Cluster 6), astrocytes (Clusters 1, 5, 26, 31, 34), oligodendrocytes (Clusters 0, 14, 20, 22, 33), oligodendrocyte precursor cells (OPCs, Clusters 8, 19), microglia (Clusters 12, 13, 24, 25, 29), pericytes (Clusters 7, 17, 18, 23), endothelium (Clusters 2, 9, 11, 21), and macrophages (Cluster 15), with comparable representation in both groups. **b**, Heatmap of log2 fold changes between No Ext and Ext by cell types. Each perpendicular line represents a differentially expressed gene (DEG).

**Extended Data Fig 2.**
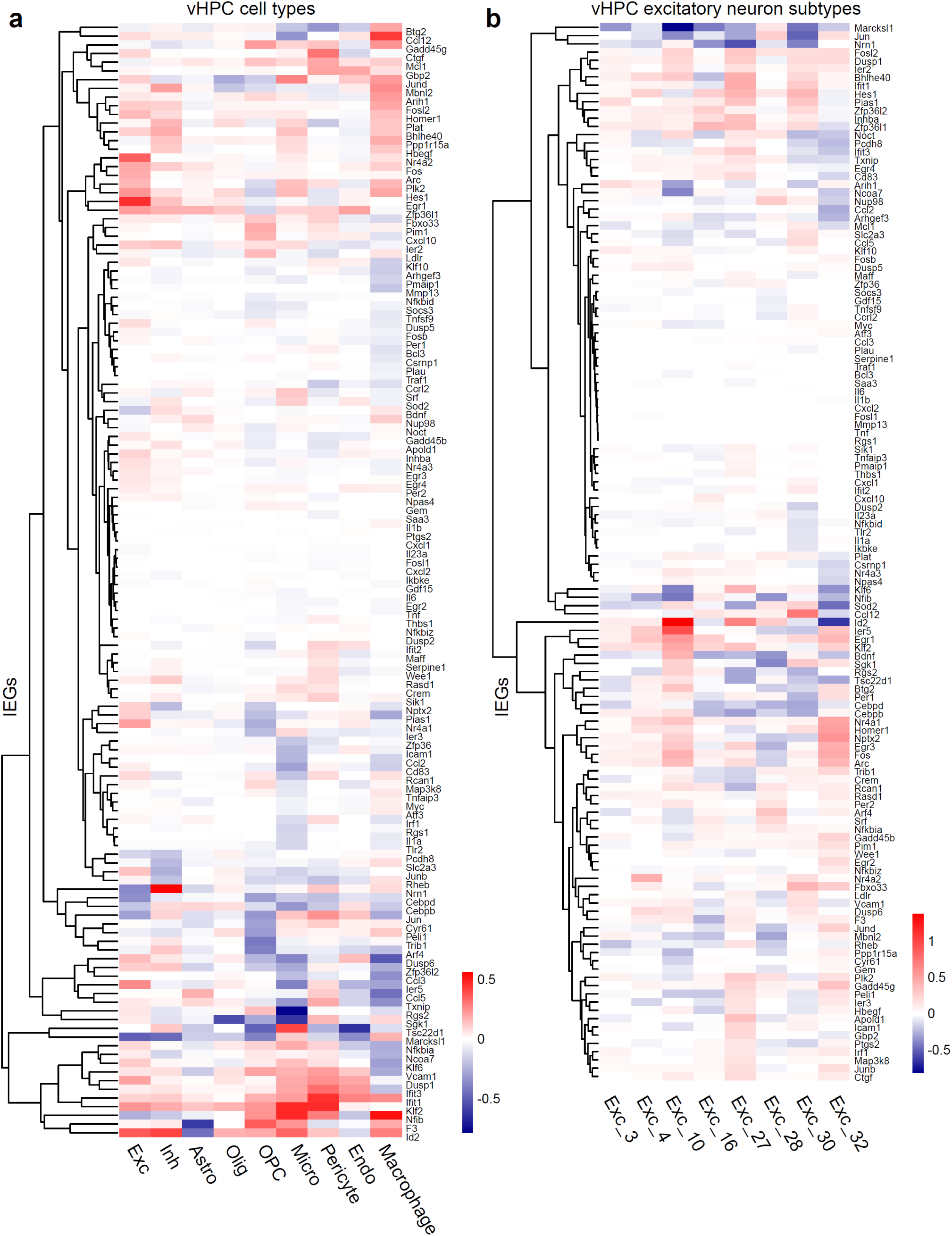
Different expression levels of IEGs across different cell types and clusters of No Ext and Ext mice. **a, b**, A clustered heatmap displaying the Log_2_ fold change profiles of the known IEGs from comparing No Ext *vs.* Ext across different cell types (**a**) and different subtypes of excitatory neurons (**b**) (columns).

**Extended Data Fig 3.**
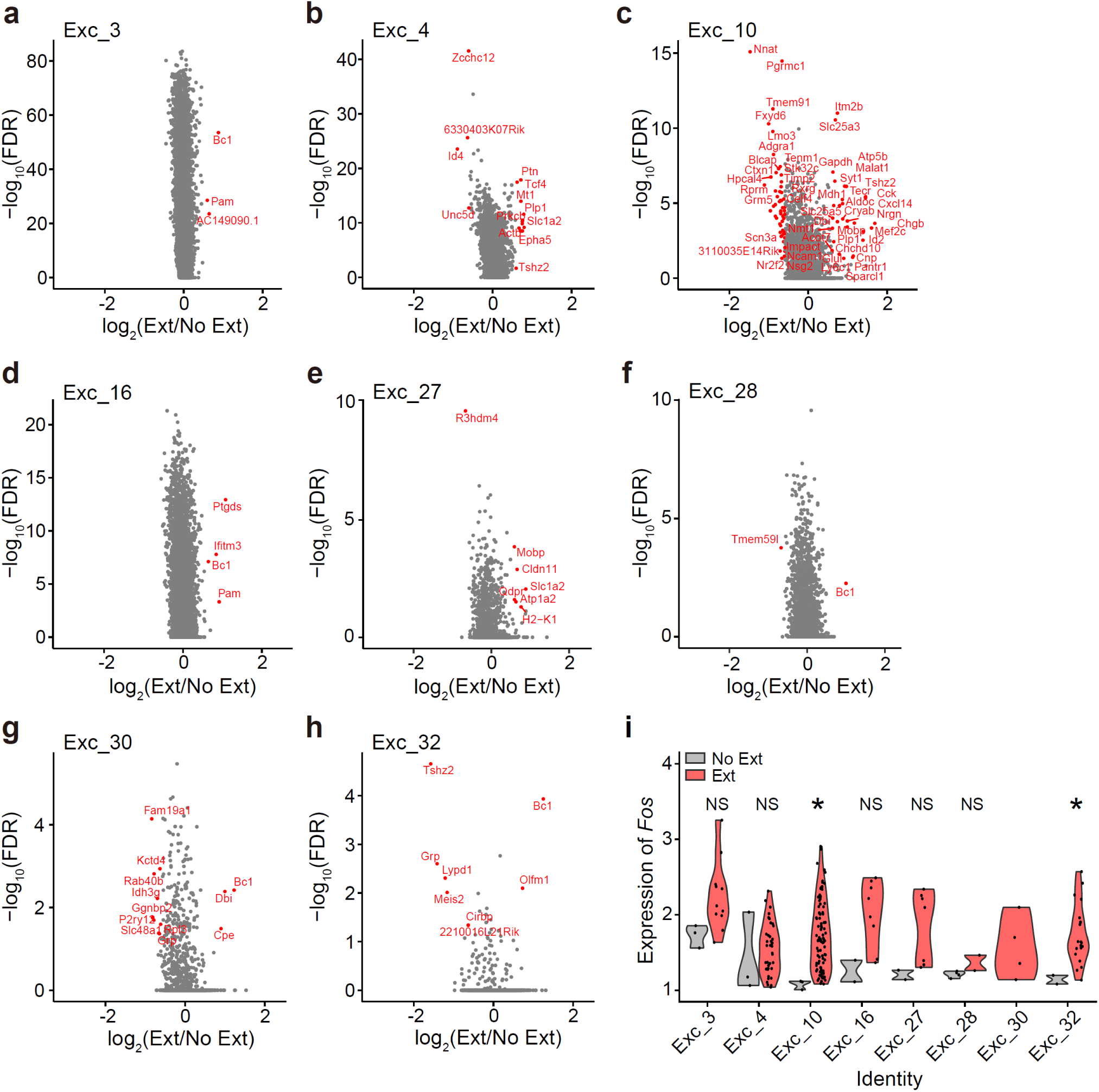
Extinction triggered varied transcriptomic program changes in all the 8 clusters. **a–h**, Volcano plots of DEGs (determined by MAST test) that were upregulated and downregulated by extinction in eight different excitatory neuron subtypes. DEGs (FDR < 0.05, |log_2_(Ext/No Ext)| > 0.585) were indicated by red dots. **i**, Violin plot of expression of *Fos* across different excitatory neuron subtypes in No Ext and Ext. NS, *P* > 0.05; Exc_10, \**P* = 0.017; Exc_22, \**P* = 0.045, unpaired Student’s *t*-test.

**Extended Data Fig 4.**
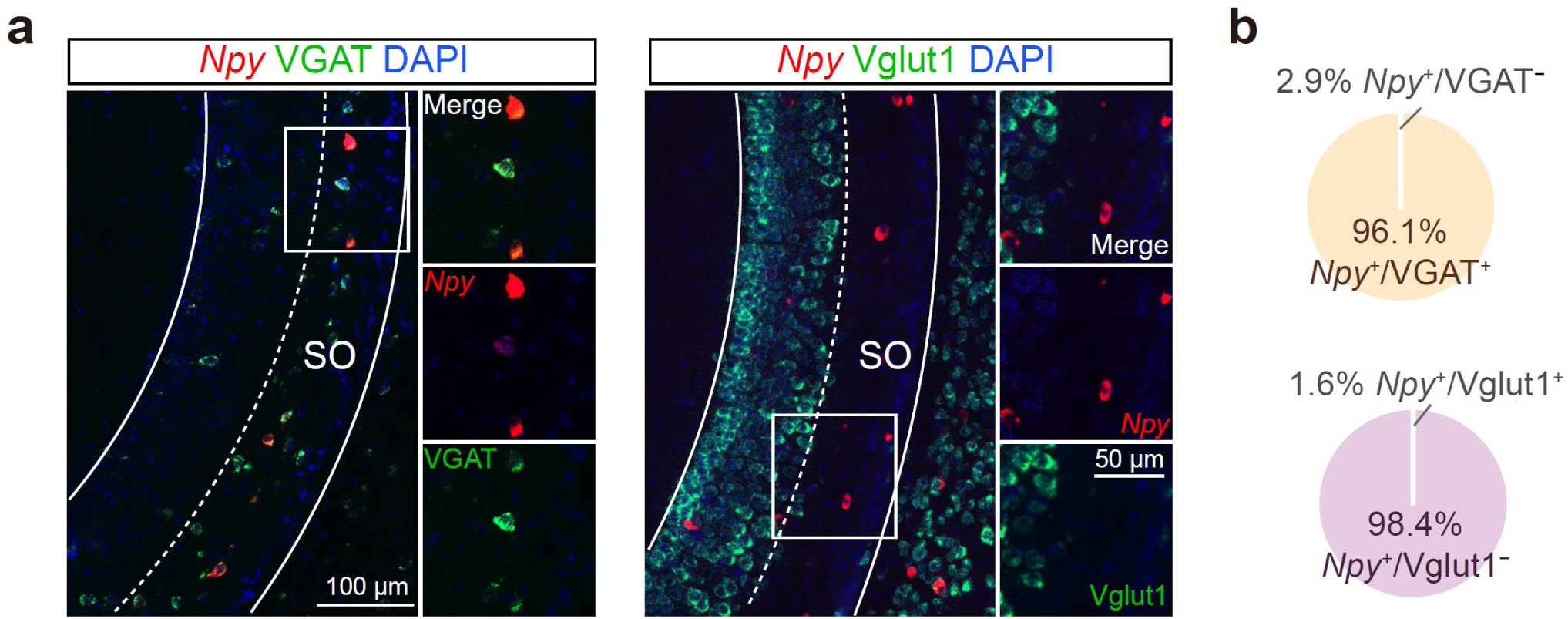
The expression of *Npy* was exclusive to VGAT^+^ neurons rather than Vglut1^+^ neurons. **a**, Dual-color FISH (dFISH) for *Npy* and GABAergic (vesicular GABA transporter, VGAT) or glutamatergic (vesicular glutamate transporter 1, Vglut1) markers differentially expressed in vCA1. SO, stratum oriens. **b**, Quantification of the percentage overlap between *Npy^+^* and molecular markers of neuronal types.

**Extended Data Fig 5.**
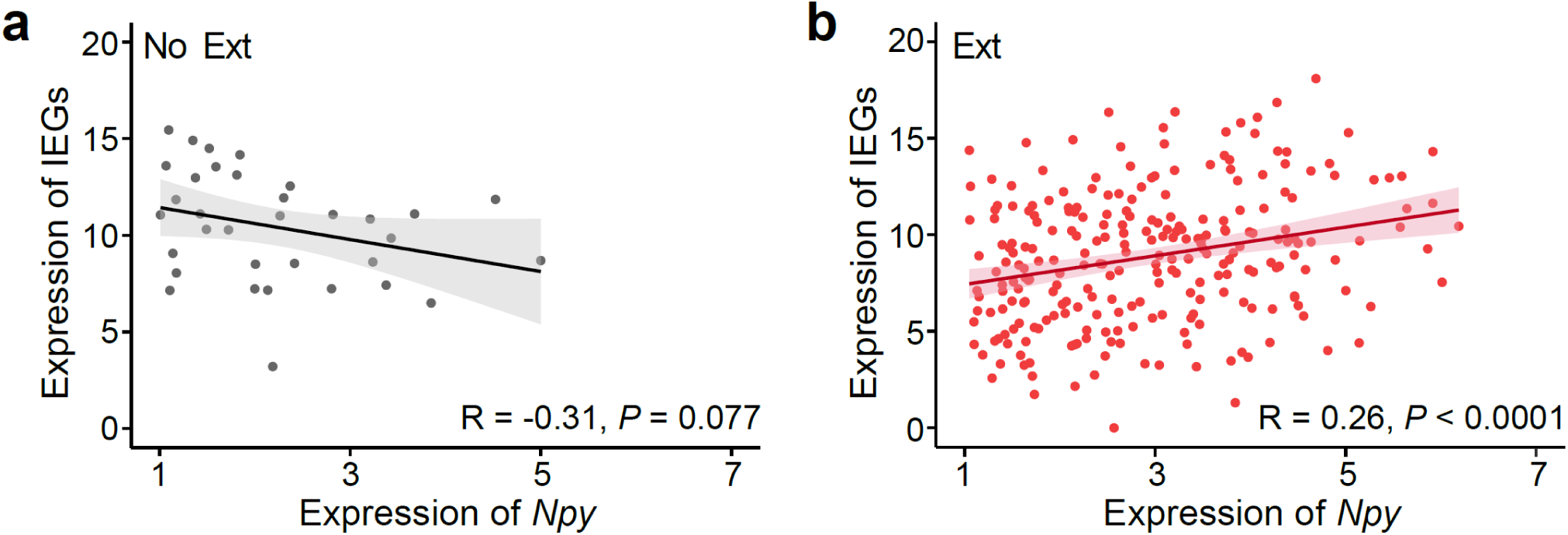
Correlation between the expression level of IEGs and *Npy* in No Ext and Ext mice. Pearson correlation coefficients and *P*-values are shown for *Npy* expression and the total expression level of 37 selected IEGs in No Ext (**a**) and Ext groups (**b**).

**Extended Data Fig 6.**
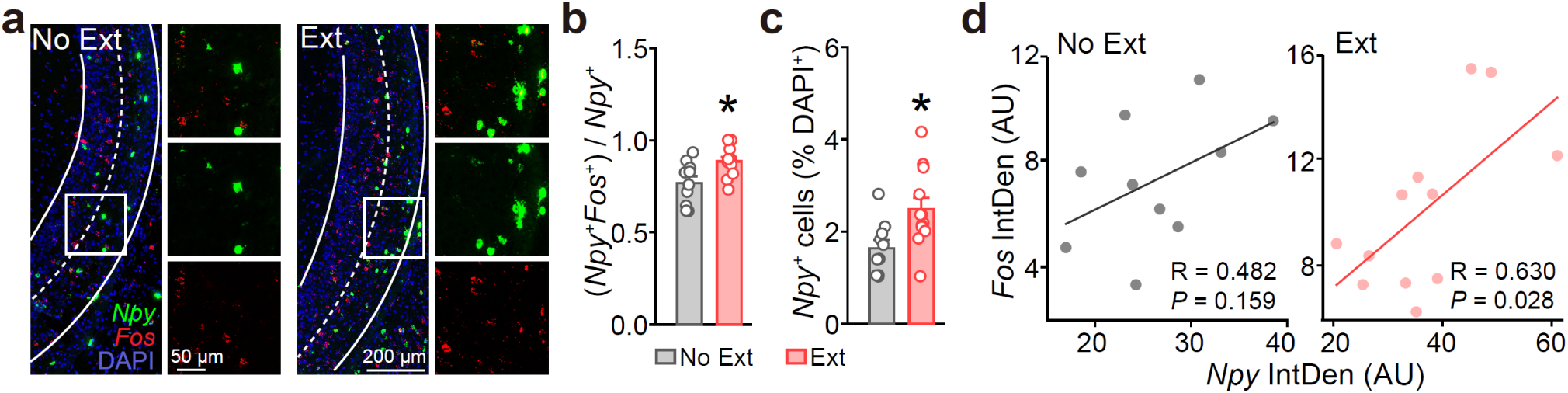
The expression level of *Npy* and *Fos* in the vCA1 exhibited a significant correlation in the Ext group but not the No Ext group. **a**, dFISH for *Npy* (green) and *Fos* (red) expression in vCA1 of No Ext and Ext mice. **b**, **c**, The proportion of activated *Npy*^+^ neurons (*Npy*^+^*/Fos*^+^, **b**) and total *Npy*^+^ neurons (*Npy*^+^, **c**) were both increased in vCA1. \**P* = 0.011 (**b**), \**P* = 0.013 (**c**), unpaired Student’s *t*-test. **d**, Pearson correlation between *Npy* and *Fos* expression levels in vCA1 of No Ext and Ext. No Ext, n = 10 brain slices of four mice, Ext, n = 12 brain slices of four mice. Correlation coefficients (R) and *P*-values (*P*).

**Extended Data Fig 7.**
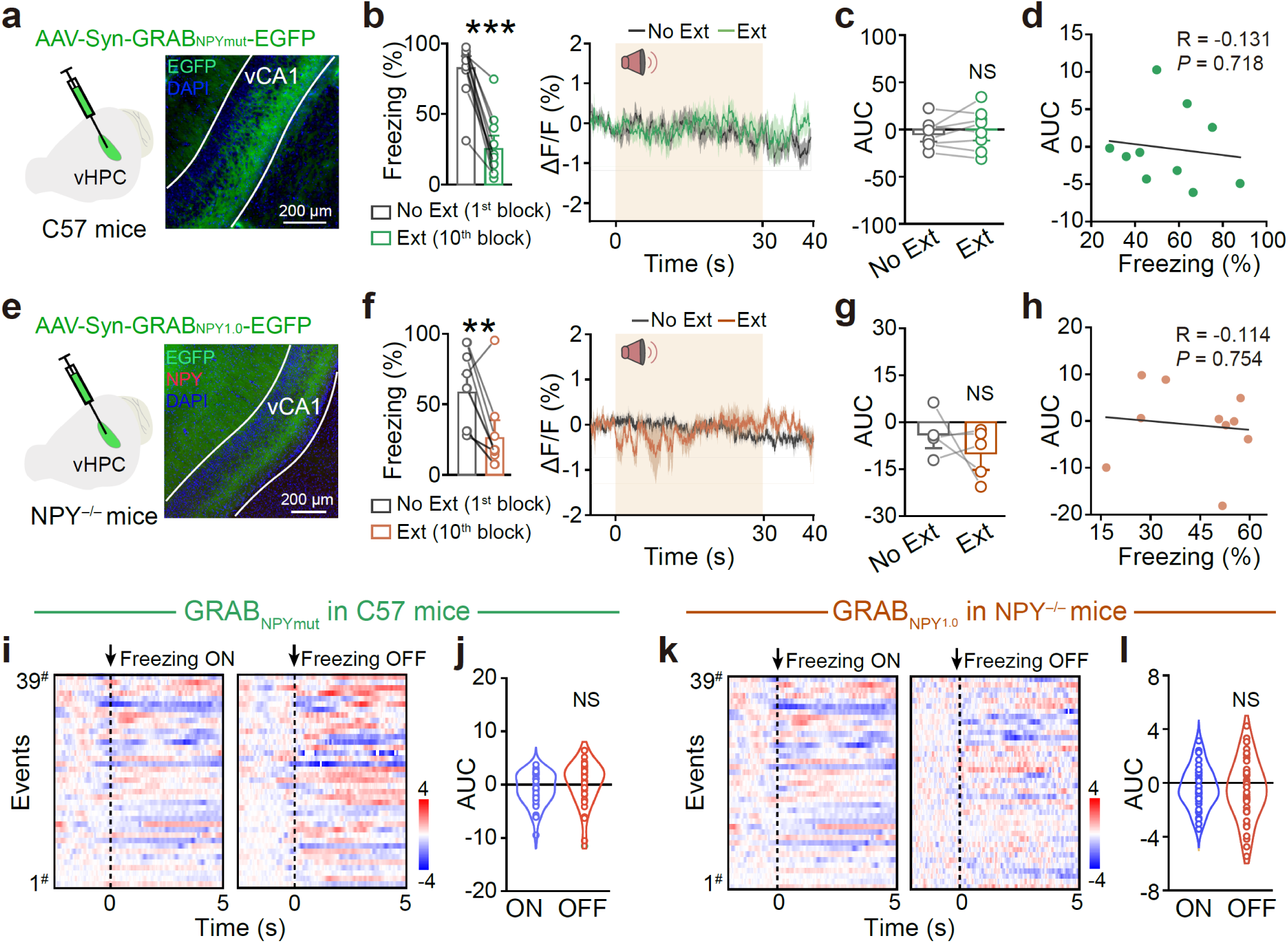
Neither GRAB_NPYmut_ sensor in wild-type mice nor the GRAB_NPY1.0_ sensor in *Npy* knockout mice exhibited any detectable response in vCA1 during extinction learning. **a**, **e**, Schematic of viral injection and expression of a genetically encoded GRAB_NPYmut_ (**a**) and GRAB_NPY1.0_ (**e**) for fiber photometry recording. **b–d**, GRAB_NPYmut_ sensor in wild-type mice did not show significant responses during extinction. n = 8 mice for No Ext, 8 mice for Ext. (**b**) Freezing levels, \*\*\**P* < 0.0001, two-tailed paired Student’s *t*-test; (**c**) NS, *P* = 0.423, two-tailed paired Student’s *t*-test; (**d**) Pearson Corr., R = –0.046, NS, *P* = 0.901. **f–h**, GRAB_NPY1.0_ sensor in NPY knockout (NPY^−/−^) mice did not show a significant response during extinction. n = 5 mice for No Ext, 5 mice for Ext. (**f**) Freezing levels, \*\**P* = 0.006, two-tailed paired Student’s *t*-test; (**g**) NS, *P* = 0.365, two-tailed paired Student’s *t*-test; (**h**) Pearson Corr., R = –0.114, NS, *P* = 0.754. **i, j**, Heatmaps (**i**) and AUC (**j**) of GRAB_NPYmut_ fluorescence during freezing ON and OFF epochs. n = 8 mice and 39 trials for freezing ON and OFF, respectively. NS, *P* = 0.224, unpaired Student’s *t*-test. **k, l**, Heatmaps (**k**) and AUC (**l**) of GRAB_NPY1.0_ fluorescence of NPY^−/−^ mice during freezing ON and OFF epochs. n = 5 mice and 39 trials for freezing ON and OFF, respectively. NS, *P* = 0.224, unpaired Student’s *t*-test.

**Extended Data Fig 8.**
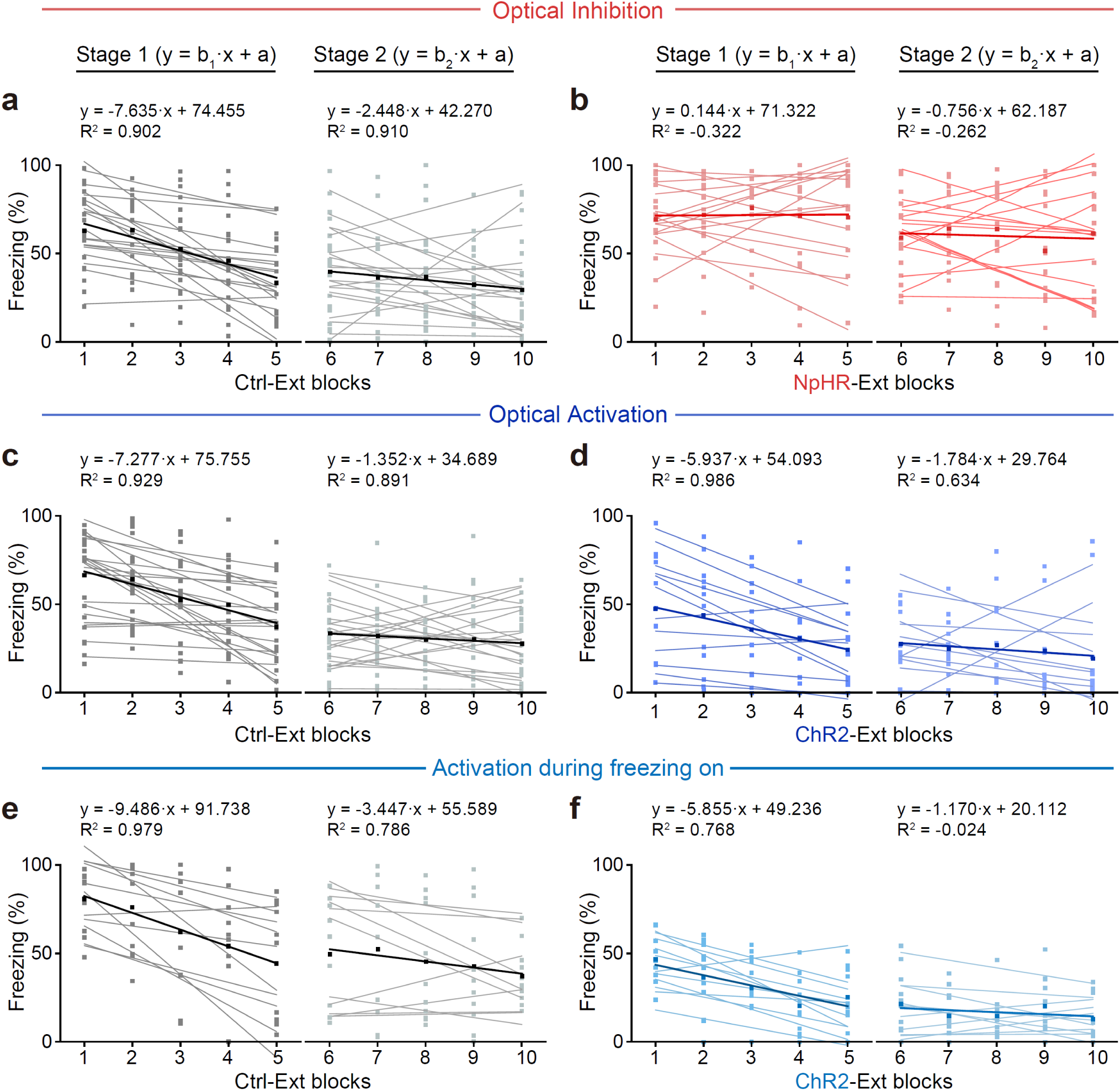
Optogenetic manipulations of NPY^+^ neuronal activity in the vCA1 altered the memory encoding during extinction. Linear fitting was performed for extinction Stage 1 and Stage 2 in mice used for optogenetic inhibition of NPY^+^ interneurons. Means are shown as thick lines. The equation and adjusted R-squared (R²) were shown for the means.

**Extended Data Fig 9.**
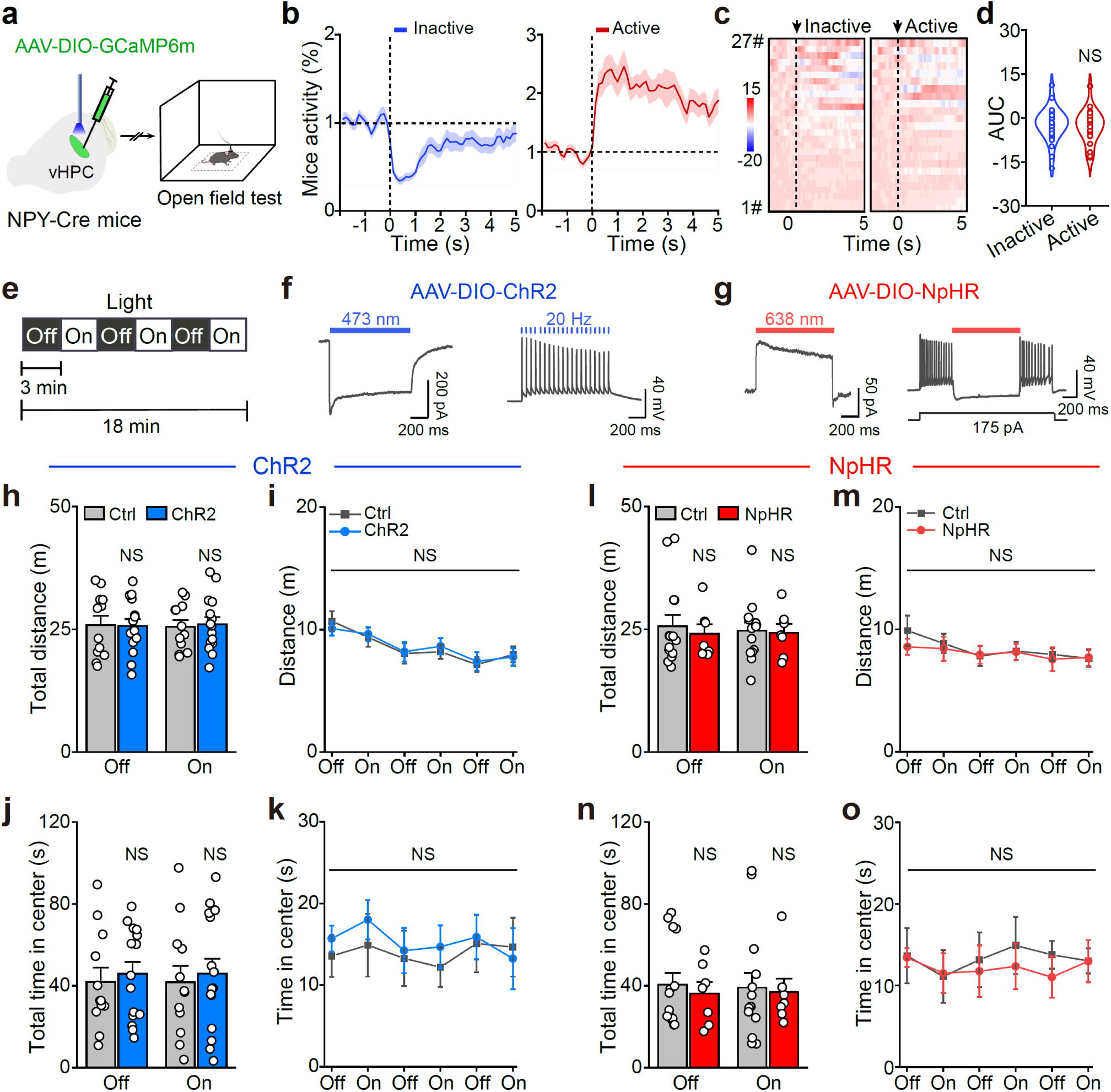
No significant correlation was found between locomotor activity and vCA1 NPY+ neuronal activity. **a**, Schematic of viral injection and the behavioral procedure for fiber photometry recording in open field tests. **b**, Based on locomotor activity, behaviors were categorized as either inactive or active in open field tests. **c**, Heatmaps of calcium responses of vCA1 NPY^+^ neurons for the corresponding inactive and active state. **d**, AUC of the ΔF/F GCaMP6m fluorescence for 5 s following locomotor activity, whether inactive or active. n = 27 trials for low motion, n = 23 trials for high motion of four mice. NS, *P* = 0.715, unpaired Student’s *t*-test. **e**, Schematic procedure for optogenetic manipulations of NPY^+^ neuronal activity in the vCA1. **f, g**, Functional identification of ChR2 (**f**) and NpHR (**g**) expressed in vCA1 NPY^+^ neurons. **h–o**, Activation or inhibition of vCA1 NPY^+^ neurons didn’t affect locomotor activity and time spent in the center. n = 14 mice for Ctrl of inhibition, n = 7 mice for NpHR, n = 12 mice for Ctrl of activation, n = 15 mice for ChR2. NS, *P* > 0.05, unpaired Student’s *t*-test.

**Extended Data Fig 10.**
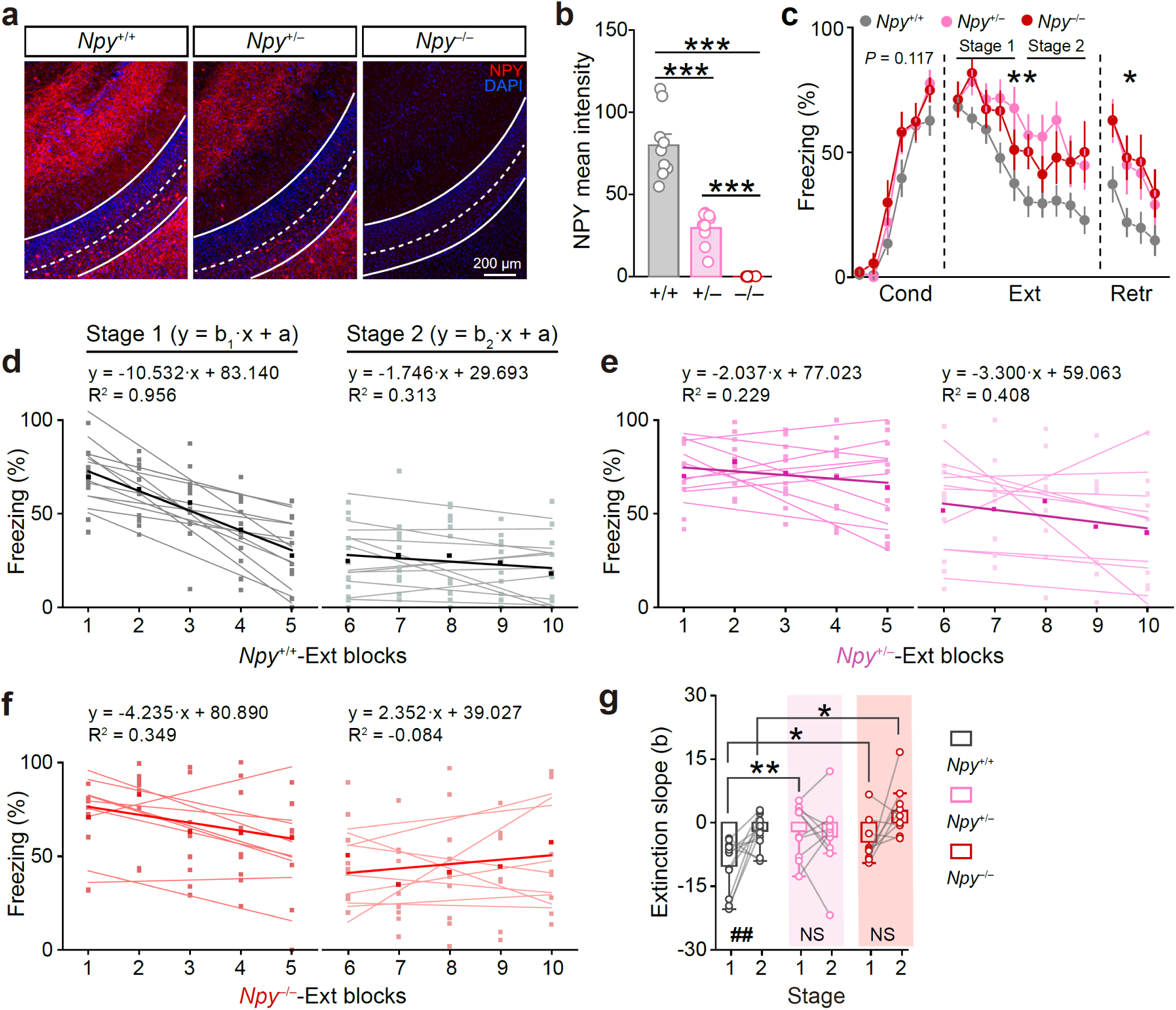
Knockdown or knockout of *Npy* impaired fear extinction. **a**, Representative images of NPY expression in vCA1 of *Npy*^+/+^, *Npy*^+/−^ and *Npy*^−/−^ mice. **b**, Quantification of NPY expression levels by IHC. n = 3 mice and 9 coronal sections for *Npy*^+/+^, *Npy*^+/−^ and *Npy*^−/−^, respectively. \*\*\**P* < 0.001, unpaired Student’s *t*-test. **c**, Extinction of *Npy*^+/−^ and *Npy*^−/−^ mice was impeded. n = 15 mice for *Npy*^+/+^, n = 9 mice for *Npy*^+/−^, n = 9 mice for *Npy*^−/−^. Two-way repeated ANOVA. Cond: F_(2, 30)_ = 2.302, NS, *P* = 0.117; *Npy*^+/+^ *vs*. *Npy*^+/−^, NS, *P* = 0.139; *Npy*^+/+^ *vs*. *Npy*^−/−^, NS, *P* = 0.058; *Npy*^+/−^ *vs*. *Npy*^−/−^, NS, *P* = 0.689. Ext: F_(2, 30)_ = 7.238, \*\**P* = 0.003; *Npy*^+/+^ *vs*. *Npy*^+/−^, \*\**P* = 0.001; *Npy*^+/+^ *vs*. *Npy*^−/−^, \**P* = 0.014; *Npy*^+/−^ *vs*. *Npy*^−/−^, NS, *P* = 0.420. Retr: F_(2, 30)_ = 4.301, \**P* = 0.023; *Npy*^+/+^ *vs*. *Npy*^+/−^, \**P* = 0.031; *Npy*^+/+^ *vs*. *Npy*^−/−^, \**P* = 0.015; *Npy*^+/−^ *vs*. *Npy*^−/−^, NS, *P* = 0.772. **d**–**f**, Linear fitting was performed for extinction Stage 1 and Stage 2 in mice used for *Npy*^+/+^, *Npy*^+/−^ and *Npy*^−/−^ mice. Means are shown as thick lines. The equation and adjusted R-squared (R²) were shown for the means. **g**, In *Npy* knockout or knockdown mice, the difference in extinction slope between Stage 1 and 2 during extinction training was lost. Stage 1 *vs.* Stage 2: *Npy*^+/+^, ^##^*P* = 0.005; *Npy*^+/−^, NS, *P* = 0.631; *Npy*^−/−^, NS, *P* = 0.100, two-tailed paired Student’s *t*-test. *Npy*^+/+^ *vs. Npy*^+/−^, \*\**P* = 0.004; *Npy*^+/+^ *vs. Npy*^−/−^, Stage 1, \**P* = 0.035, Stage 2, \**P* = 0.045; unpaired Student’s *t*-test.

**Extended Data Fig 11.**
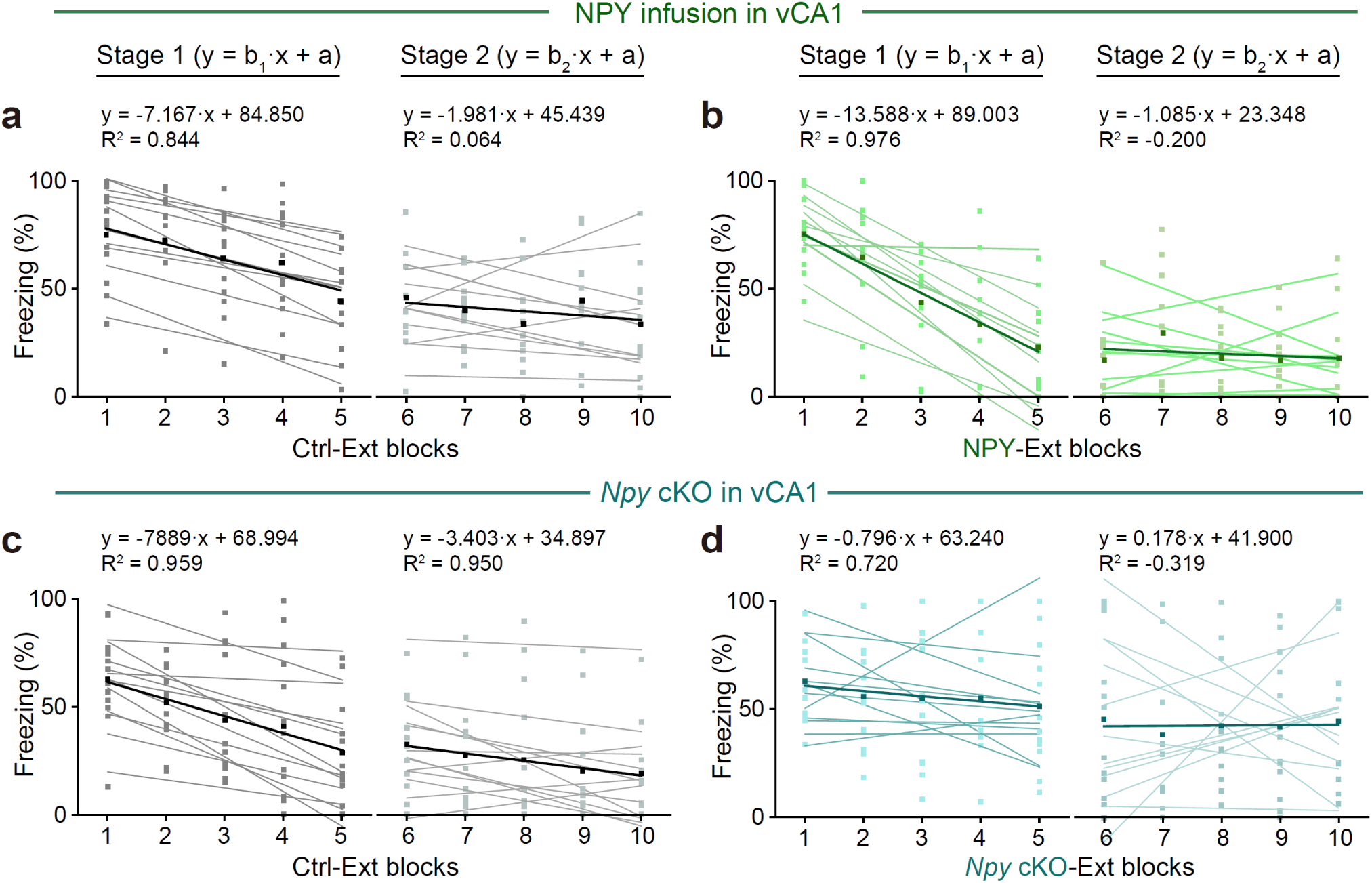
NPY itself is both necessary and sufficient for fear extinction. Linear fitting was performed for extinction Stage 1 and Stage 2 in mice used for pharmacology or genetic manipulation NPY levels. Means are shown as thick lines. The equation and adjusted R-squared (R²) were shown for the means.

**Extended Data Fig 12.**
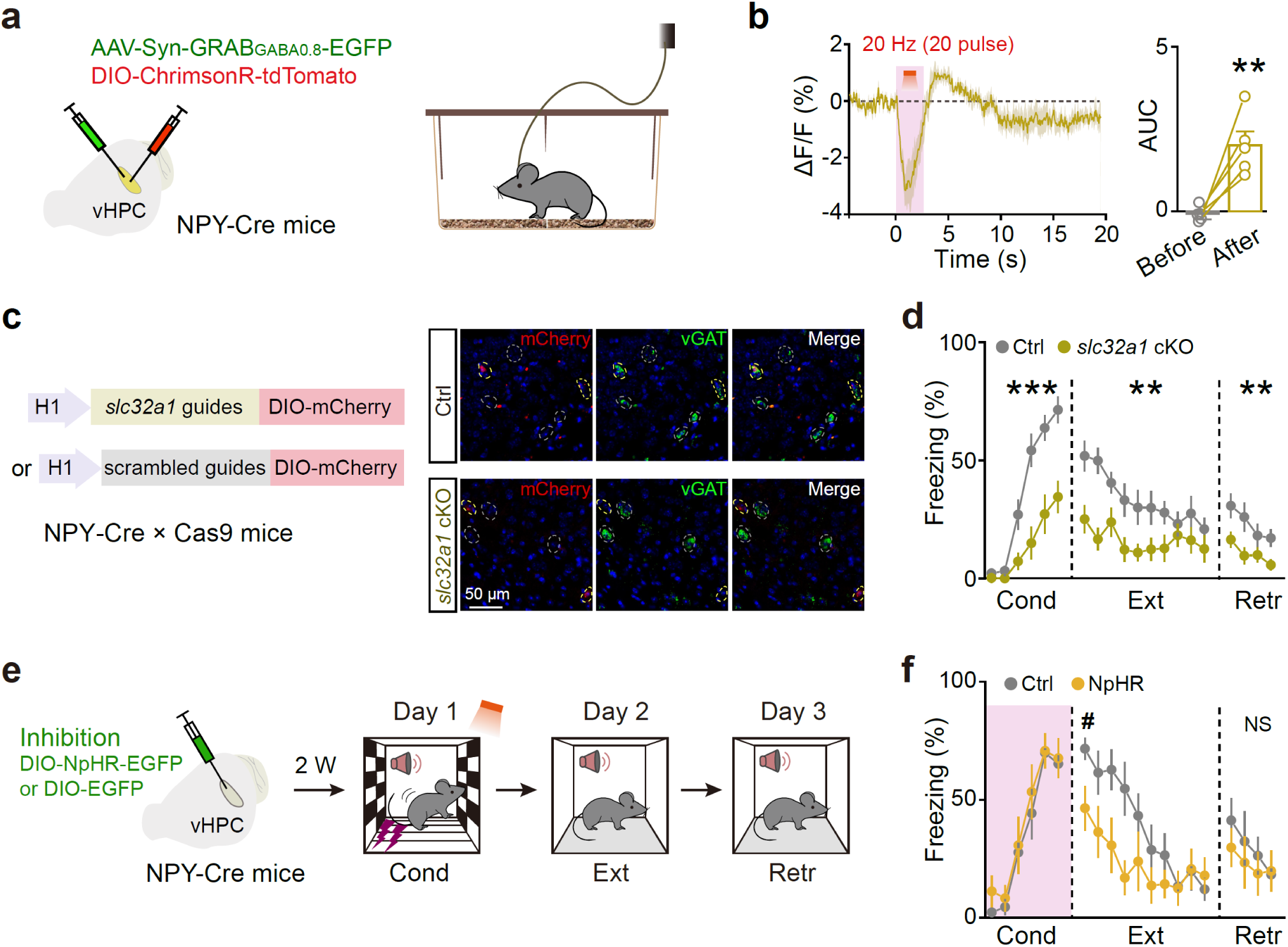
Both *slc32a1* cKO in the vCA1 NPY+ interneurons, and inhibiting NPY^+^ neuronal activity during conditioning, significantly weakened fear memory. **a**, Schematic of viral injection for fiber photometry. **b**, GRAB_GABA0.8_ sensors responses before (5 s) and after stimulating NPY^+^ neurons. n = 5 mice. \*\**P* = 0.009, two-tailed paired Student’s *t*-test. **c**, Schematic of *slc32a1* cKO in the vCA1 NPY^+^ neurons and images showing the expression of vGAT and mCherry in the control and *slc32a1* cKO groups. Yellow circles with mCherry expression represent virus-infected neurons, gray circles represent vGAT-expressing neurons. In the *slc32a1* cKO group, there was virtually no vGAT expression in virus-infected neurons compared to non-virally infected neurons in the same slice or to the neurons in the Ctrl group with or without virus infection. **d**, *slc32a1* cKO significantly impeded fear conditioning. n = 14 and 14 mice for Ctrl and *slc32a1* cKO. Two-way repeated ANOVA, main effects of group. Cond: F_(1, 26)_ = 23.355, \*\*\**P* < 0.001. Ext: F_(1, 26)_ = 12.376, \*\**P* = 0.002. Retr: F_(1, 26)_ = 8.417, \*\**P* = 0.007. **e**, Schematics of viral injection and behavioral procedure for optogenetic inhibition of NPY^+^ neurons in the vCA1. **f**, Inhibition of NPY^+^ neurons during conditioning decreased fear memory retrieval. n = 8 mice for Ctrl and 7 mice for NpHR. Two-way repeated ANOVA, main effects of group, Ext: F_(1, 13)_ = 4.060, NS, *P* = 0.065. Retr: F_(1, 13)_ = 0.430, NS, *P* = 0.523. **^#^***P* = 0.022, unpaired Student’s *t*-test.

**Extended Data Fig 13.**
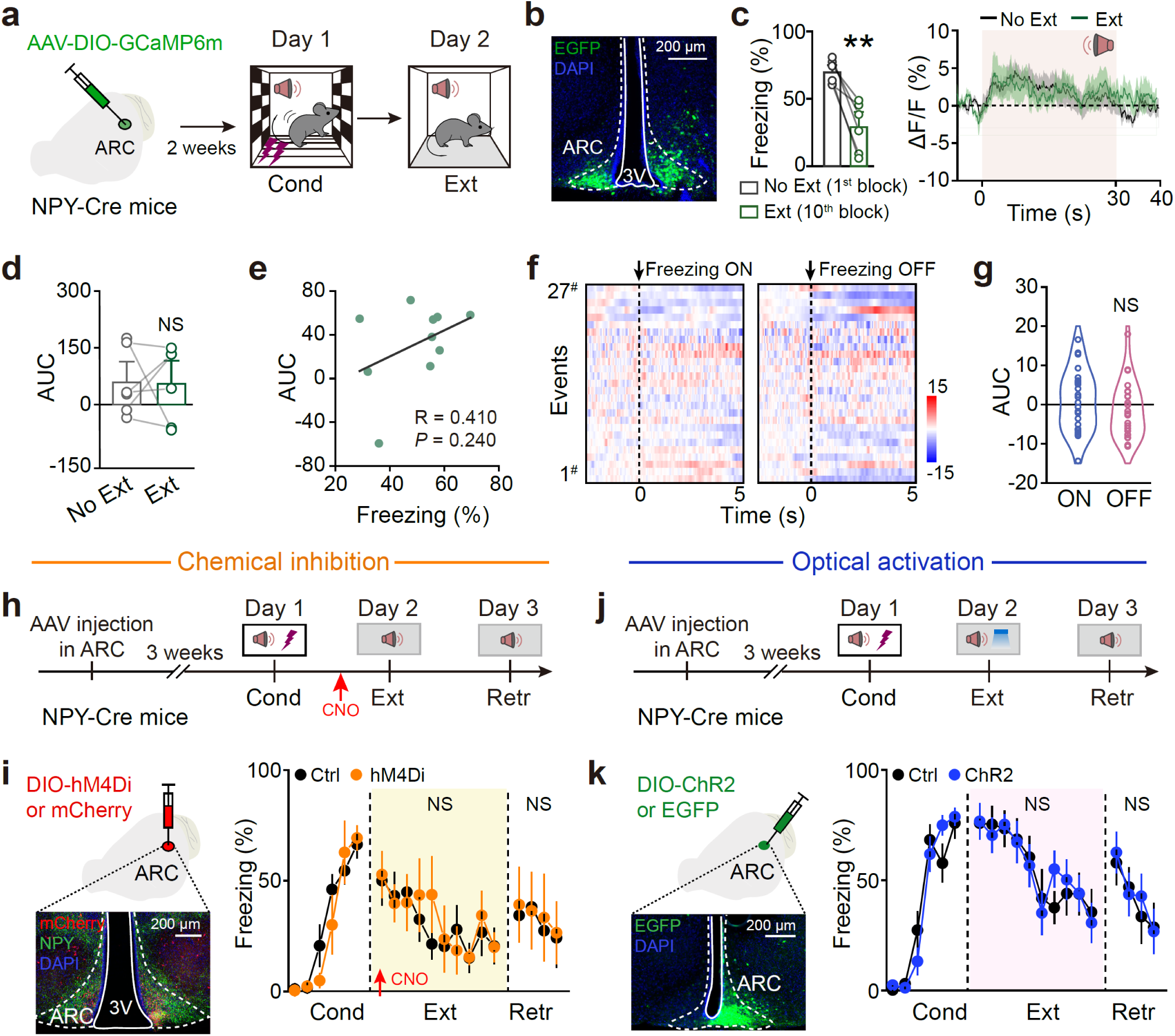
NPY^+^ neurons in the hypothalamic arcuate nucleus have no impact on fear memory. **a**, Schematic of viral injection and the behavioral procedure for fiber photometry recording in ARC. **b**, Representative image showing the expression of GCaMP6m in ARC. **c–e**, The response of GCaMP6m in the ARC did not have significant difference between the Ext and No Ext state. n = 6 mice. (**c**) Freezing levels, **\*\****P* = 0.006, two-tailed paired Student’s *t*-test; (**d**) NS, *P* = 0.953, two-tailed paired Student’s *t*-test; (**e**) Pearson Corr., R = 0.410, *P* = 0.240. **f, g**, Heatmaps (**f**) and AUC (**g**) of GCaMP6m responses during freezing ON and OFF. n = 6 mice and 27 trials for freezing ON and OFF, respectively. NS, *P* = 0.454, unpaired Student’s *t*-test. **h, j**, Schematic behavioral procedure for chemical inhibition and optical activation of NPY^+^ neurons in the ARC. **i**, Left: Representative image showing the expression of hM4Di-mCherry in ARC NPY^+^ neurons; Right: Chemical inhibition of NPY^+^ neurons in ARC had no effect on extinction. n = 7 mice for Ctrl, and 5– 6 mice for hM4Di. Two-way repeated ANOVA. Ext: F_(1, 11)_ = 0.088, NS, *P* = 0.772. Retr: F_(1, 10)_ = 0.031, NS, *P* = 0.863. **k**, Left: Representative image showing the expression of ChR2-EGFP in ARC NPY^+^ neurons. Right: Activation of NPY^+^ neurons in ARC failed to affect fear extinction. n = 8 mice for Ctrl, and 10 mice for ChR2. Two-way repeated ANOVA. Ext: F_(1, 16)_ = 0.005, NS, *P* = 0.945. Retr: F_(1, 16)_ = 0.038, NS, *P* = 0.848.

**Extended Data Fig 14.**
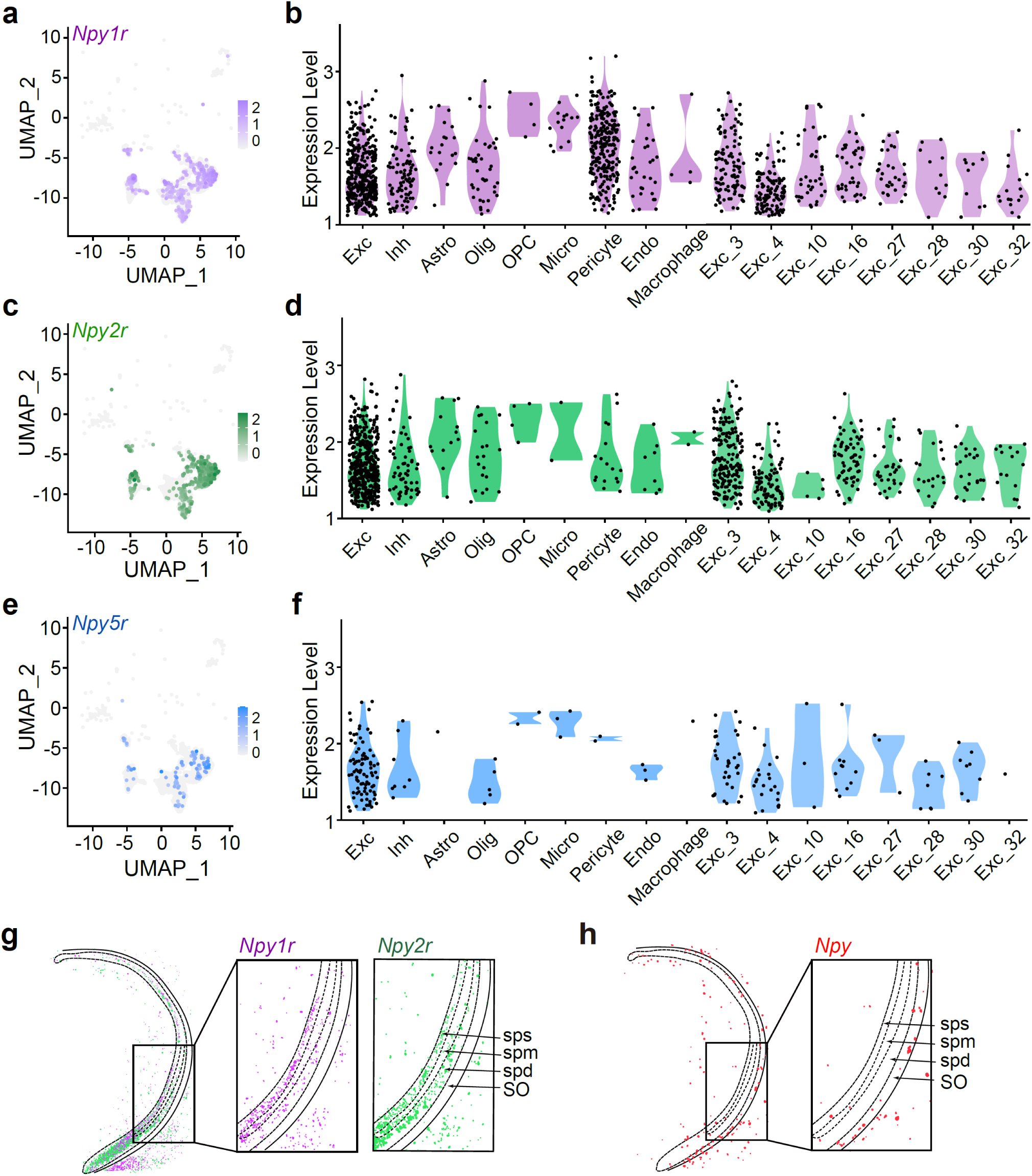
Expression and distribution of NPY receptors in vHPC. **a, c, e**, UMAP representation of *Npy1r^+^, Npy2r^+^ and Npy5r^+^*clusters in vHPC. **b, d, f**, Violin plots showing the expression levels of *Npy1r^+^, Npy2r^+^ and Npy5r^+^* across different cell types and excitatory neuron subtypes in vHPC. Each dot represents a single cell. **g**, **h**, RNAscope images depicting the regional and laminar specificities of *Npy1r*, *Npy2r* and *Npy* gene expression in vHPC. vCA1-sps, superficial pyramidal layer of vCA1; vCA1-spm, middle pyramidal layer of vCA1; vCA1-spd, deep pyramidal layer of vCA1.

**Extended Data Fig 15.**
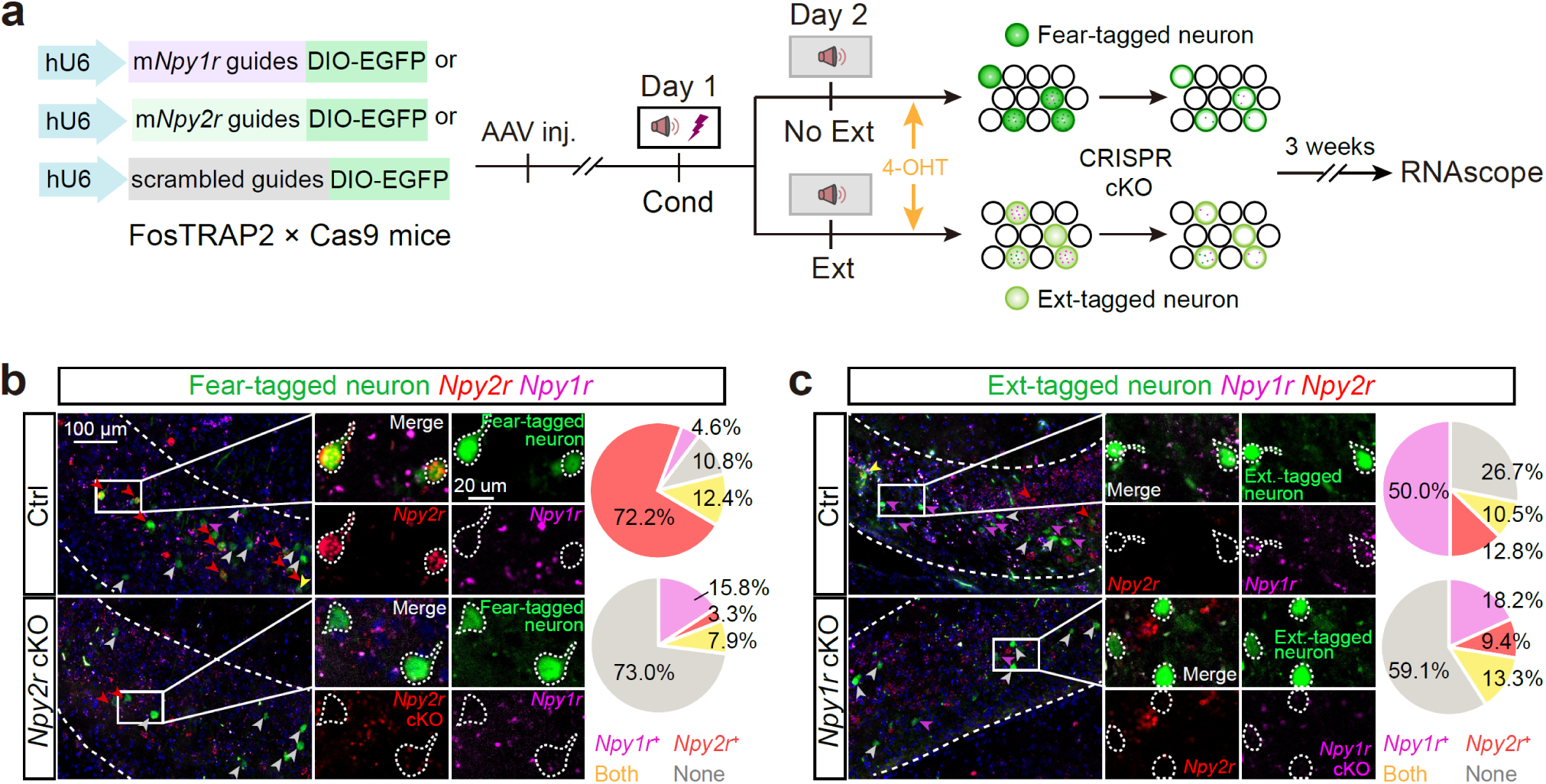
*In vivo* validation of CRISPR/Cas9-mediated conditioned KO (cKO) of *Npy2r* or *Npy1r* in Fear- and Ext-tagged neurons. **a**, Experimental design to validate the efficacy of the CRISPR/Cas9 system in depleting *Npy2r* and *Npy1r* mRNA levels in Fear- and Ext-tagged neurons, respectively. **b**, Images from a representative brain slice showing the expression of *Npy2r*, *Npy1r* in the Fear-tagged and *Npy2r* cKO neurons. The pie chart indicates that after *Npy2r* cKO in Fear-tagged neurons, the relative proportion of *Npy2r*-expressing neurons was significantly decreased. n = 12 brain slice of three mice for Ctrl; n = 17 brain slice of two mice for *Npy2r* cKO. **c**, Images from a representative brain slice showing *Npy1r*, *Npy2r* in the Ext-tagged and *Npy1r* cKO neurons. The pie chart indicates that after *Npy1r* cKO in Ext-tagged neurons, the relative proportion of *Npy1r*-expressing neurons was significantly decreased. n = 10 brain slice of two mice for Ctrl, n = 31 brain slice of three mice for *Npy1r* cKO.

**Extended Data Fig 16.**
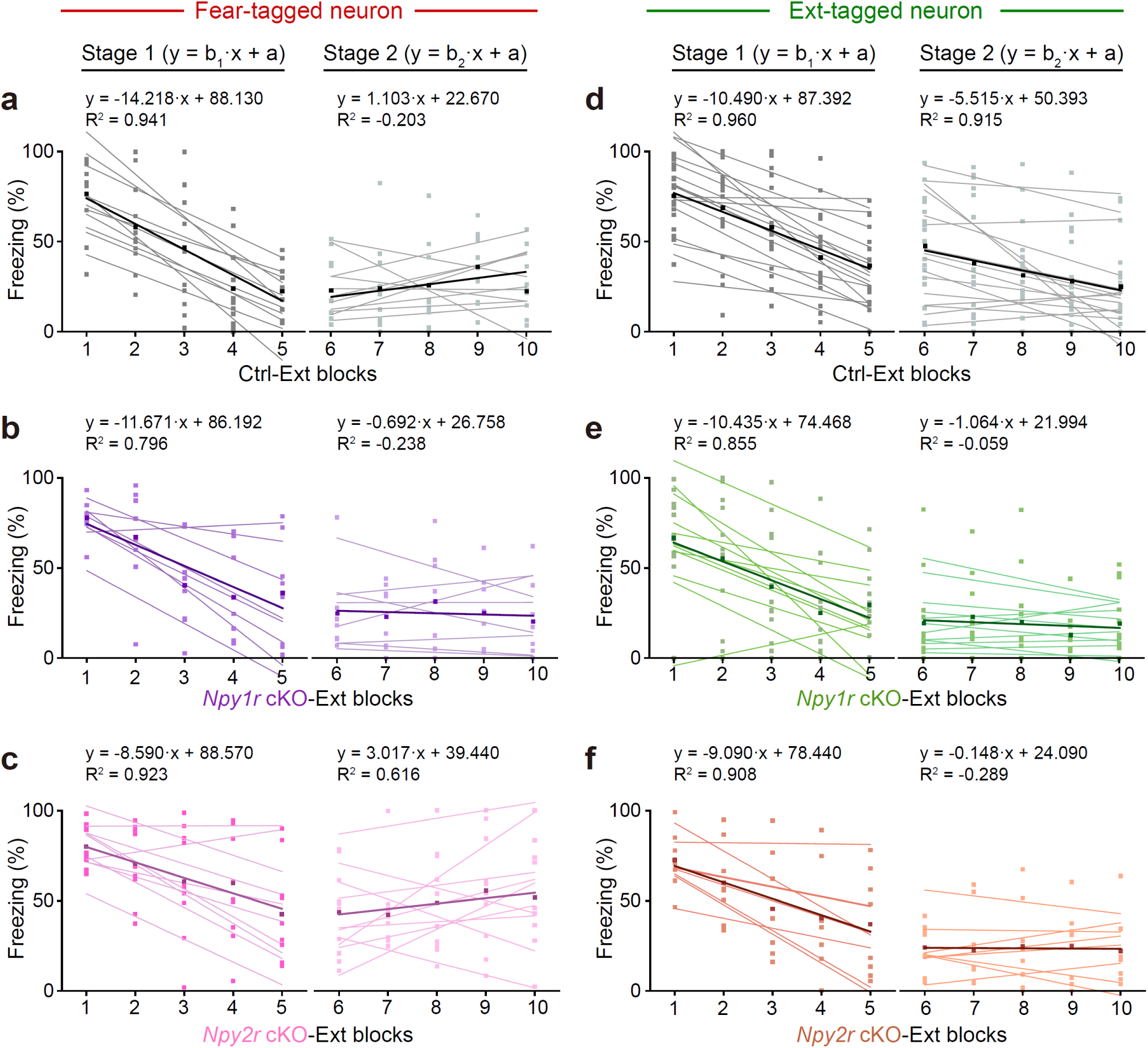
*Npy* receptor gene cKO in fear- or Ext-tagged neurons altered fear and extinction memory. Linear fitting was performed for extinction Stage 1 and Stage 2 in mice used for *Npy* receptor gene cKO. Means are shown as thick lines. The equation and adjusted R-squared (R²) were shown for the means.

**Extended Data Fig 17.**
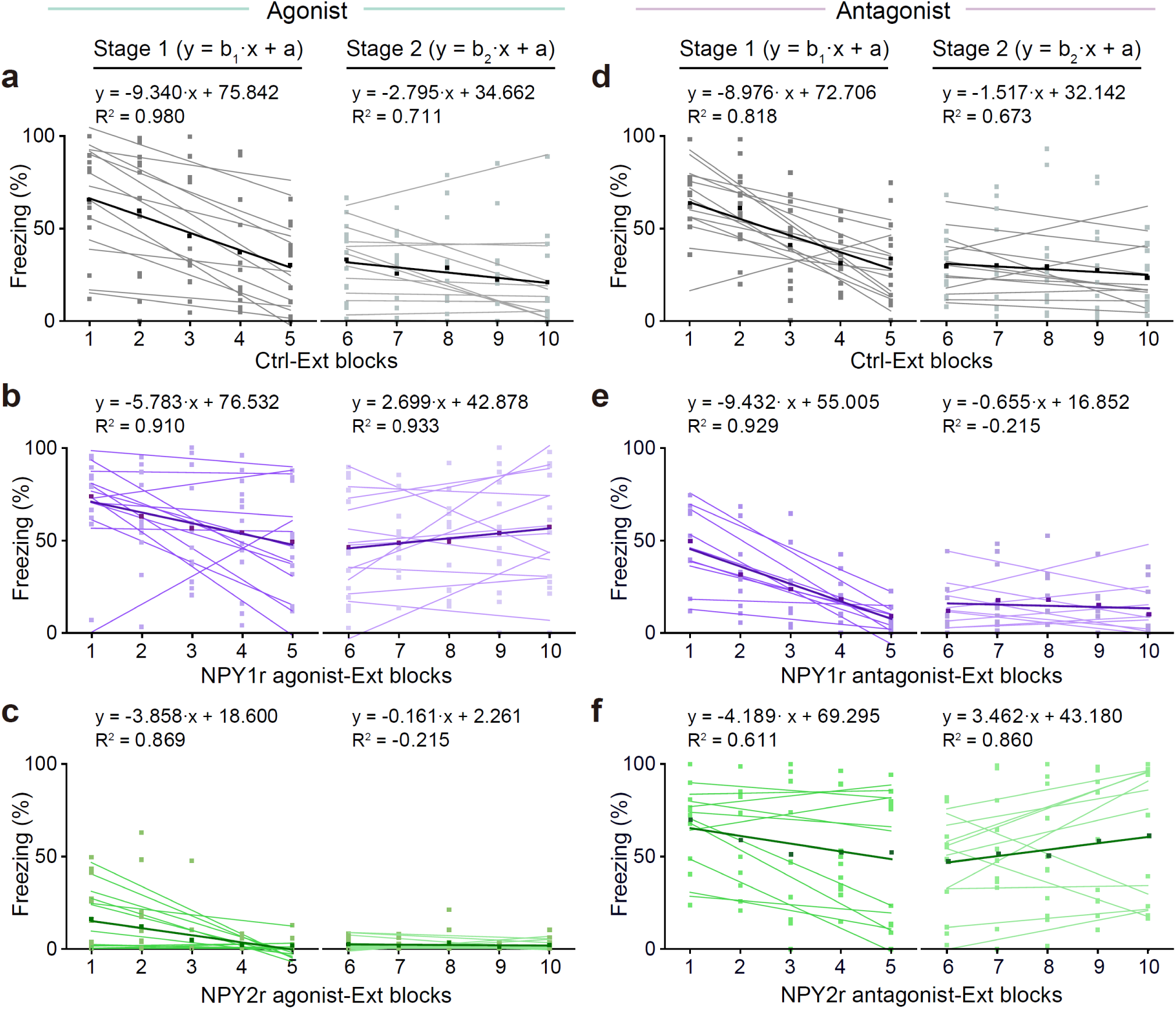
Pharmacological modulation of NPY1r and NPY2r activity impacted fear and extinction memory. Linear fitting was performed for extinction Stage 1 and Stage 2 in mice used for pharmacological modulation of NPY receptor activity. Means are shown as thick lines. The equation and adjusted R-squared (R²) were shown for the means.

**Extended Data Fig 18.**
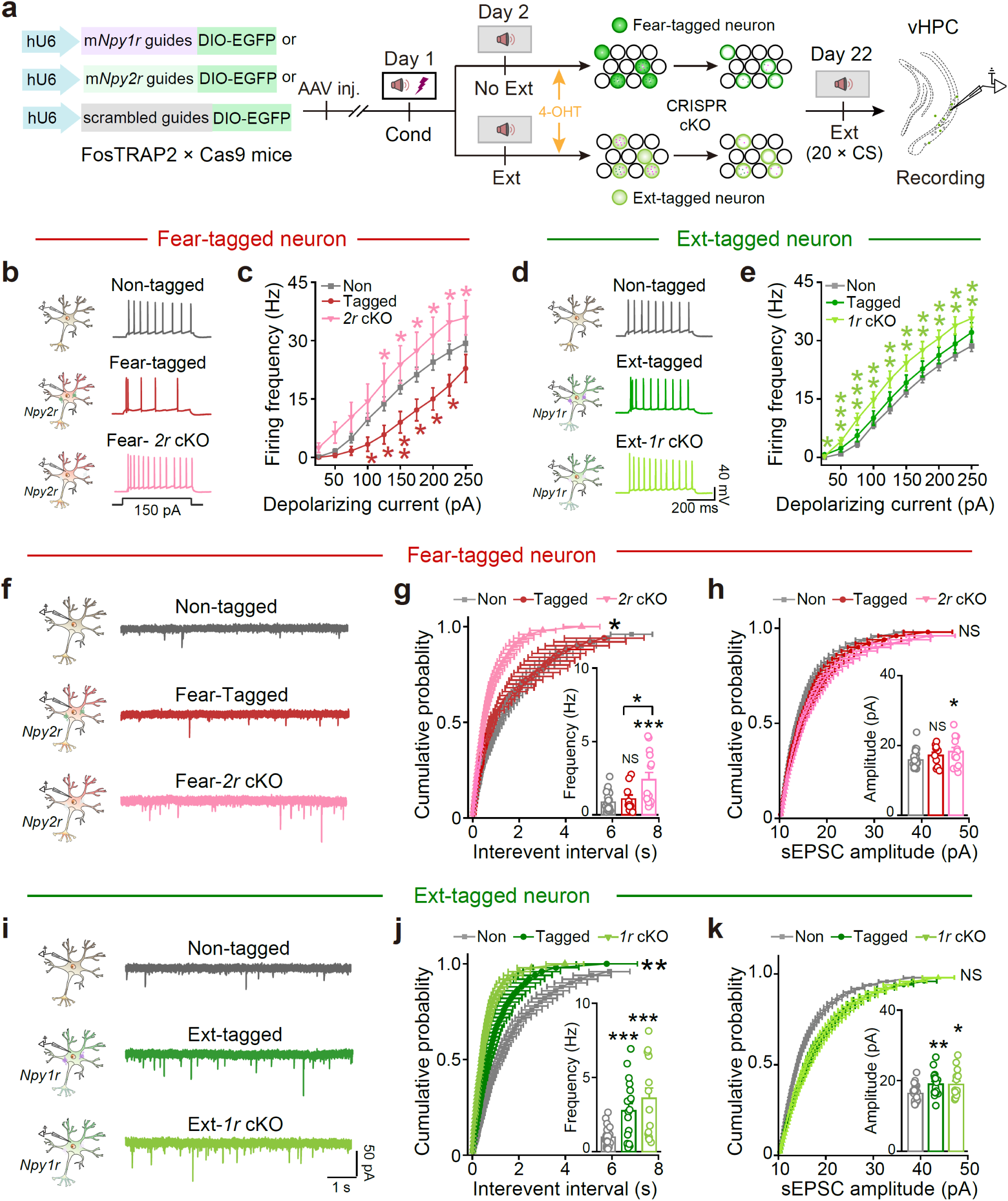
Following extinction, recording the action potentials and sEPSCs of fear-tagged and Ext-tagged neurons with or without Npy receptor gene cKO. Following extinction, fear-tagged neurons exhibited significant decreases in AP and sEPSC frequencies, which were reversed by *Npy2r* KO selectively in these neurons. By contrast, the Ext-tagged neurons showed no significant increase in AP frequencies but exhibited increased sEPSC frequencies. *Npy1r* KO in the Ext-tagged neurons increased their AP firing frequencies. **a**, Behavioral procedure for electrophysiological recordings, and schematic of *Npy2r*, *Npy1r* gene cKO in Fear- and Ext-tagged neurons respectively. **b**, Representative traces of AP firing evoked by current injection (150 pA) in Non-, Fear-tagged and *Npy2r* cKO neurons. **c**, *Npy2r* cKO in Fear-tagged neurons significantly reversed the decreased firing frequency following extinction. n = 26 cells of eight mice for Non-tagged, n = 12 cells of four mice for Fear-tagged, n = 17 cells of five mice for *Npy2r* cKO. \**P* < 0.05, \*\**P* < 0.01, unpaired Student’s *t*-test, Non-tagged *vs. Npy2r* cKO (red asterisk), Fear-tagged *vs. Npy2r* cKO (pink asterisk). **d**, Representative traces of AP firing evoked by current injection (150 pA) in Non-, Ext-tagged and *Npy1r* KO neurons. **e**, *Npy1r* cKO in Ext-tagged neurons enhanced the firing frequency compared to Non-tagged neurons after extinction. n = 43 cells of thirteen mice for Non-tagged, n = 19 cells of four mice for Fear-tagged, n = 34 cells of six mice for *Npy1r* cKO. \**P* < 0.05, \*\**P* < 0.01, \*\*\**P* < 0.001, unpaired Student’s *t*-test, Non-tagged *vs. Npy1r* cKO (light green asterisk). **f**, Representative traces of sEPSCs in Non-, Fear-tagged and *Npy2r* cKO neurons following extinction. **g**, *Npy2r* cKO in Fear-tagged neurons increased the sEPSC frequency. Non-tagged *vs.* Fear-tagged, NS, *P* = 0.967; Non-tagged *vs. Npy2r* cKO, \**P* = 0.022; Fear-tagged *vs. Npy2r* cKO, NS, *P* = 0.068, two-sample Kolmogorov-Smirnov test; Non-tagged *vs.* Fear-tagged, NS, *P* = 0.448; Non-tagged *vs. Npy2r* cKO, \*\*\**P* < 0.001; Fear-tagged *vs. Npy2r* cKO, \**P* = 0.045, unpaired Student’s *t*-test. **h**, *Npy2r* cKO in Fear-tagged neurons didn’t increase the sEPSC amplitude. Non-tagged *vs.* Fear-tagged, NS, *P* = 0.967; Non-tagged *vs. Npy2r* cKO, NS, *P* = 0.869; Fear-tagged *vs. Npy2r* cKO, NS, *P* = 1.000, two-sample Kolmogorov-Smirnov test; Non-tagged *vs.* Fear-tagged, NS, *P* = 0.191; Non-tagged *vs. Npy2r* cKO, \**P* = 0.024; Fear-tagged *vs. Npy2r* cKO, NS, *P* = 0.470, unpaired Student’s t-test. n = 26 cells of ten mice for Non-tagged, n = 10 cells of four mice for Fear-tagged, n = 14 cells of six mice for *Npy2r* cKO. **i**, Representative traces of sEPSCs in Non-, Ext-tagged and *Npy1r* cKO neurons following extinction. **j**, *Npy1r* cKO in Ext-tagged neurons increased the sEPSC frequency. Non-tagged *vs.* Ext-tagged, NS, *P* = 0.179; Non-tagged *vs. Npy1r* cKO, \*\**P* = 0.001; Ext-tagged *vs. Npy1r* cKO, NS, *P* = 0.179, two-sample Kolmogorov-Smirnov test; Non-tagged *vs.* Ext-tagged, \*\*\**P* < 0.001; Non-tagged *vs. Npy1r* cKO, \*\*\**P* < 0.001; Ext-tagged *vs. Npy1r* cKO, NS, *P* = 0.320, unpaired Student’s t-test. **k,** *Npy1r* cKO in Ext-tagged neurons didn’t increase the sEPSC amplitude. Non-tagged *vs.* Ext-tagged, NS, *P* = 0.549; Non-tagged *vs. Npy1r* cKO, NS, *P* = 0.549; Ext-tagged *vs. Npy1r* cKO, NS, *P* = 1.000, two-sample Kolmogorov-Smirnov test; Non-tagged *vs.* Ext-tagged, \*\**P* = 0.004; Non-tagged *vs. Npy1r* cKO, \**P* = 0.012; Ext-tagged *vs. Npy1r* cKO, NS, *P* = 0.941, unpaired Student’s *t*-test. n = 26 cells of eleven mice for Non-tagged, n = 18 cells of six mice for Ext-tagged, n = 15 cells of six mice for *Npy1r* cKO.

**Extended Data Fig 19.**
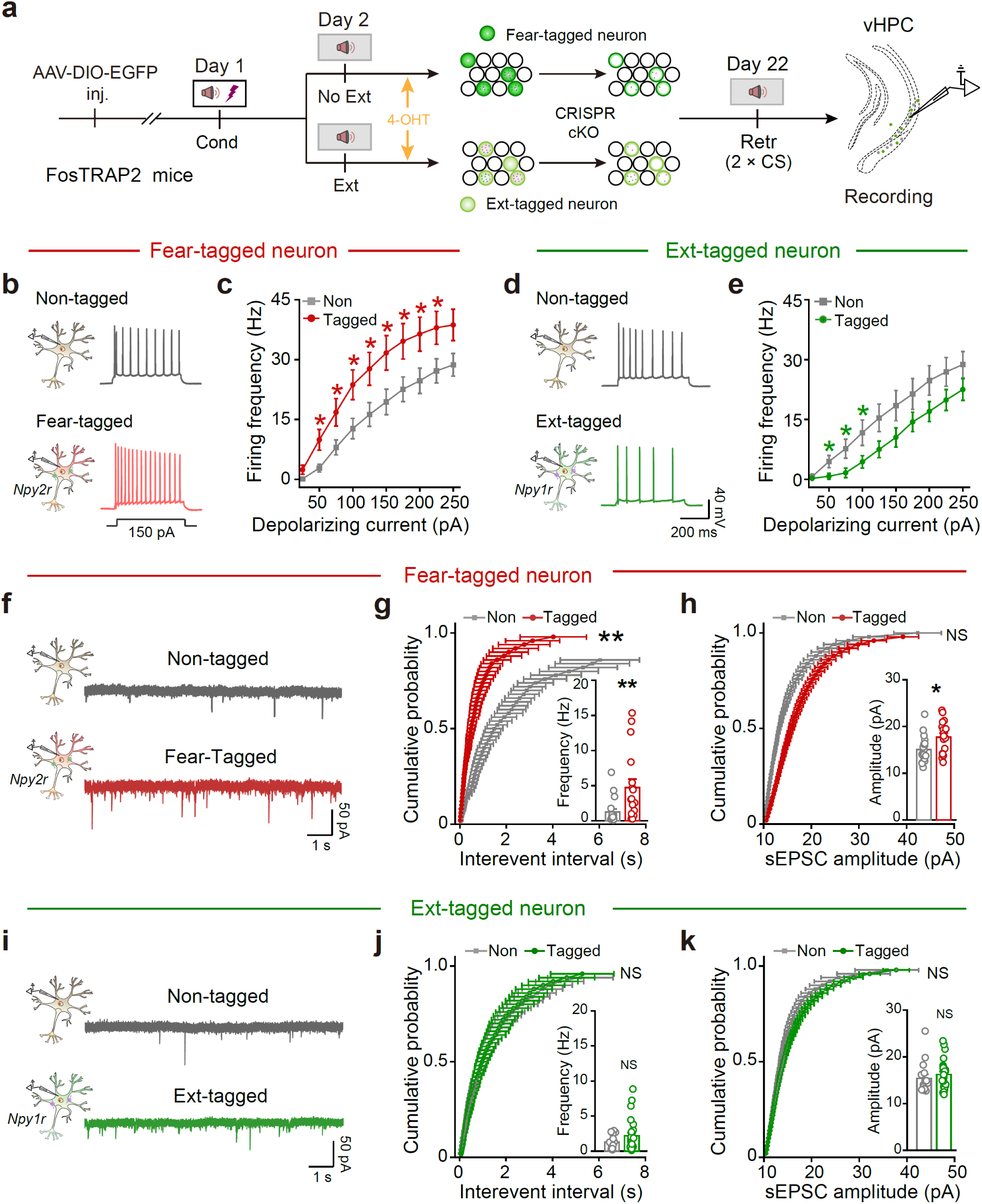
Following fear memory retrieval, recording the APs and sEPSCs of Fear- and Ext-tagged neurons. Following spontaneous recovery as fear memory retrieval with heightened fear, fear-tagged neurons displayed increased AP and sEPSC frequencies while Ext-tagged neurons exhibited decreased AP frequencies. **a**, Schematic for recording Fear- and Ext-tagged neurons following fear memory retrieval. **b**, **d**, Representative traces of AP firing evoked by current injection (150 pA) in Non-, Fear-tagged (**b**) and Ext-tagged neurons (**d**). **c**, **e**, AP frequencies of fear-tagged neurons significantly increased compared to non-tagged neurons (**c**), while Ext-tagged neurons significantly decreased (**e**), following fear memory retrieval on Day 22. (**c**) \**P* < 0.05, unpaired Student’s *t*-test, n = 17 cells of three mice for Non-tagged, n = 20 cells of three mice for Fear-tagged; (**e**) \**P* < 0.05, unpaired Student’s *t*-test, n = 15 cells of three mice for Non-tagged, n = 16 cells of three mice for Ext-tagged. **f, i**, Representative traces of sEPSCs. **g, h**, The frequency of sEPSC in Fear-tagged neurons increased, but the amplitude increased slightly, following fear memory retrieval. (**g**) \*\**P* = 0.001, two-sample Kolmogorov-Smirnov test; \*\**P* < 0.001, unpaired Student’s *t*-test. (**h**) NS, *P* = 0.112, two-sample Kolmogorov-Smirnov test; \**P* = 0.018, unpaired Student’s *t*-test. n = 18 cells of three mice for Non-tagged, n = 17 cells of three mice for Fear-tagged. **j, k**, Following spontaneous recovery as fear memory retrieval, there were no significant changes in the frequency and amplitude of sEPSCs in Ext-tagged neurons. (**j**) NS, *P* = 1.000, two-sample Kolmogorov-Smirnov test, NS, *P* = 0.172, unpaired Student’s *t*-test; (**k**) NS, *P* = 0.967, two-sample Kolmogorov-Smirnov test, NS, *P* = 0.370, unpaired Student’s *t*-test. n = 15 cells of three mice for Non-tagged, n = 24 cells of four mice for Ext-tagged.

**Extended Data Fig 20.**
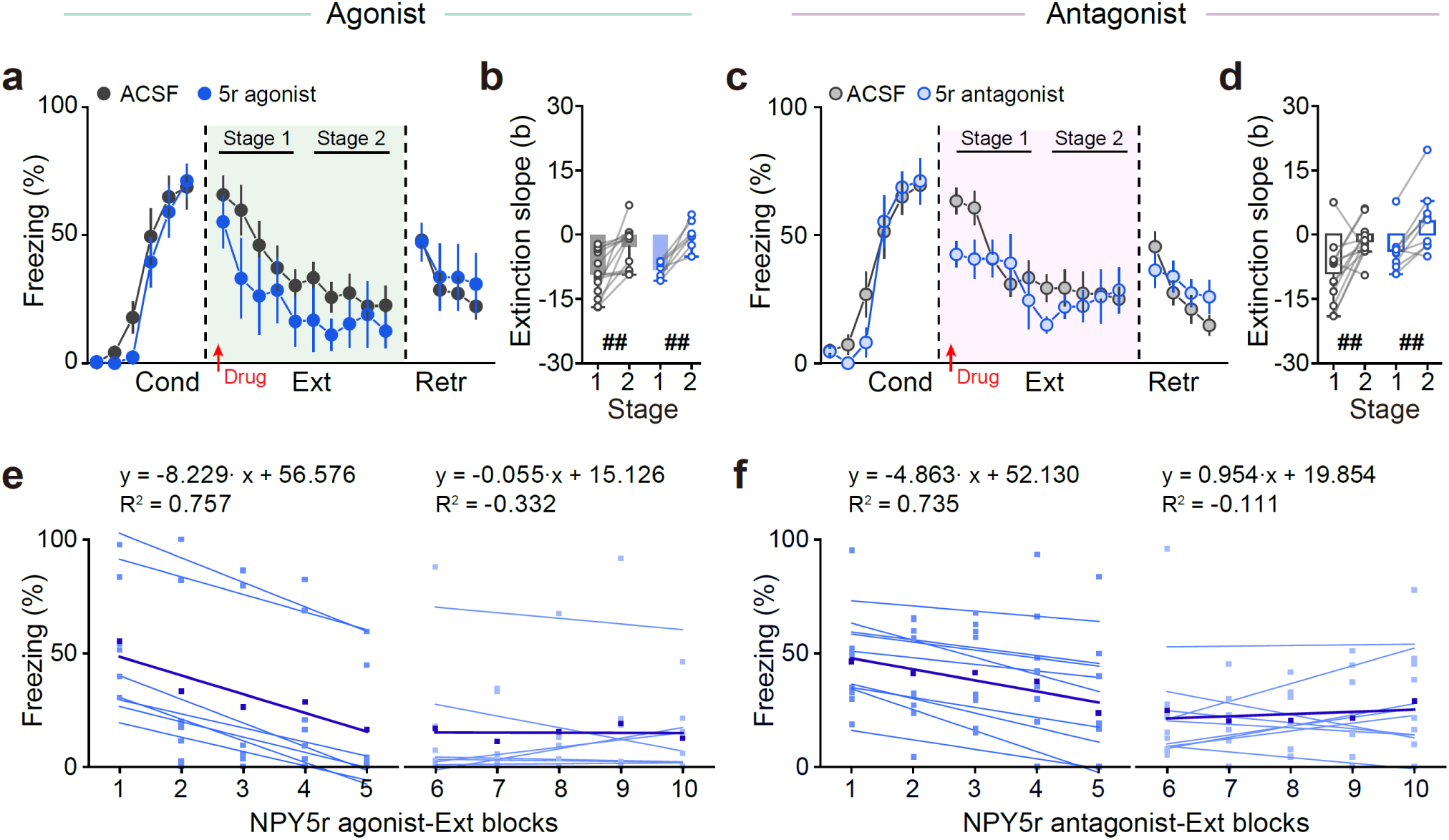
Pharmacological interventions targeting NPY5r in vCA1 had no discernible effects on fear extinction. **a–d**, Microinjection the agonist (**a, b**) and antagonist (**c, d**) of NPY5r in vCA1 had no effect on extinction. (**a**) n = 13 mice for ACSF, n = 7 mice for NPY5r agonist. Two-way repeated ANOVA, Ext: F_(1, 18)_ = 1.635, NS, *P* = 0.217; Retr: F_(1, 18)_ = 0.250, NS, *P* = 0.623. (**b**) Stage 1 *vs.* Stage 2: ACSF, ^##^*P =* 0.003; NPY5r agonist, ^##^*P* = 0.001, two-tailed paired Student’s *t*-test. (**c**) n = 13 mice for ACSF, n = 9 mice for NPY5r antagonist. Two-way repeated ANOVA. Ext: F_(1, 20)_ = 1.188, NS, *P* = 0.289; Retr: F_(1, 20)_ = 0.381, NS, *P* = 0.544. (**d**) Stage 1 *vs.* Stage 2: ACSF, ^##^*P =* 0.006; NPY5r antagonist, ^##^*P* = 0.002, two-tailed paired Student’s *t*-test. **e, f**, Linear fitting was performed for extinction Stage 1 and Stage 2 in mice with microinjection the agonist (**e**) and antagonist (**f**) of NPY5r. Means are shown as thick lines. The equation and adjusted R-squared (R²) were shown for the means. The freezing level of ACSF control group taken from Fig. 5g, h and Fig. 5i, j, respectively, is regraphed again for comparison.

**Extended Data Fig 21.**
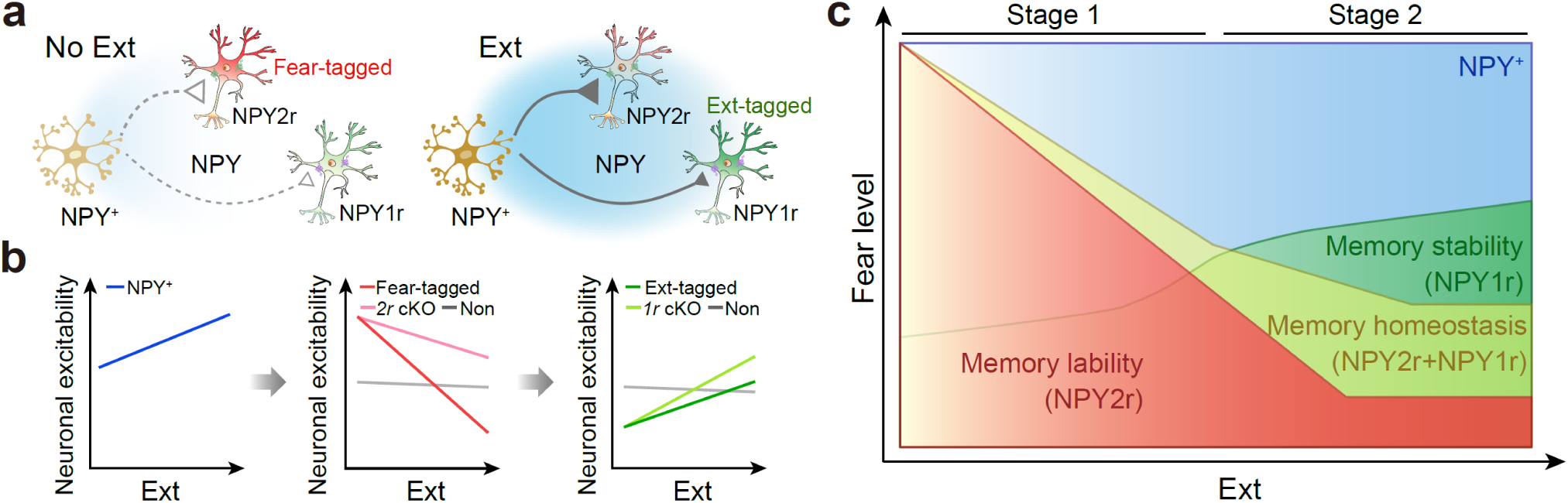
Scheme illustrating NPY modulation of neuronal ensembles for memory lability and stability. In contrast to No Ext, during extinction, NPY^+^ neuronal activity progressively increases. The increased inhibition from released NPY to *Npy2r-*expressing Fear-tagged neurons gradually intensifies, but this inhibition is negated by *Npy2r* cKO in Fear-tagged neurons. Likewise, Ext-tagged neuronal activity increases progressively during extinction. The inhibition of NPY to *Npy1r-*expressing Ext-tagged neurons is less pronounced; however, *Npy1r* cKO enhances the activity of Ext-tagged neurons. Overall, locally released NPY in the vCA1 target distinct *Npy2r-* and *Npy1r*-subensembles to establish a setpoint between memory lability and stability.

## Notes

### Competing Interest Statement

The authors have declared no competing interest.

## REFERENCES

1 Josselyn, S. A., Kohler, S. & Frankland, P. W. Finding the engram. Nat Rev Neurosci 16, 521–534 (2015). 10.1038/nrn4000

2 Josselyn, S. A. & Tonegawa, S. Memory engrams: Recalling the past and imagining the future. Science 367 (2020). 10.1126/science.aaw4325

3 Sun, X. et al. Functionally Distinct Neuronal Ensembles within the Memory Engram. Cell 181, 410–423 e417 (2020). 10.1016/j.cell.2020.02.055

4 Guskjolen, A. & Cembrowski, M. S. Engram neurons: Encoding, consolidation, retrieval, and forgetting of memory. Mol Psychiatry 28, 3207–3219 (2023). 10.1038/s41380-023-02137-5

5 Tome, D. F. et al. Dynamic and selective engrams emerge with memory consolidation. Nat Neurosci 27, 561–572 (2024). 10.1038/s41593-023-01551-w

6 Wu, Y. E., Pan, L., Zuo, Y., Li, X. & Hong, W. Detecting Activated Cell Populations Using Single-Cell RNA-Seq. Neuron 96, 313–329 e316 (2017). 10.1016/j.neuron.2017.09.026

7 Hrvatin, S. et al. Single-cell analysis of experience-dependent transcriptomic states in the mouse visual cortex. Nat Neurosci 21, 120–129 (2018). 10.1038/s41593-017-0029-5

8 Hochgerner, H. et al. Neuronal types in the mouse amygdala and their transcriptional response to fear conditioning. Nat Neurosci 26, 2237–2249 (2023). 10.1038/s41593-023-01469-3

9 Choi, J. H. et al. Interregional synaptic maps among engram cells underlie memory formation. Science 360, 430–435 (2018). 10.1126/science.aas9204

10 Gallinaro, J. V., Gasparovic, N. & Rotter, S. Homeostatic control of synaptic rewiring in recurrent networks induces the formation of stable memory engrams. PLoS Comput Biol 18, e1009836 (2022). 10.1371/journal.pcbi.1009836

11 Vogels, T. P., Sprekeler, H., Zenke, F., Clopath, C. & Gerstner, W. Inhibitory plasticity balances excitation and inhibition in sensory pathways and memory networks. Science 334, 1569–1573 (2011). 10.1126/science.1211095

12 Davis, P., Zaki, Y., Maguire, J. & Reijmers, L. G. Cellular and oscillatory substrates of fear extinction learning. Nat Neurosci 20, 1624–1633 (2017). 10.1038/nn.4651

13 Wang, Q. et al. Insular cortical circuits as an executive gateway to decipher threat or extinction memory via distinct subcortical pathways. Nat Commun 13, 5540 (2022). 10.1038/s41467-022-33241-9

14 Kim, T. et al. Activated somatostatin interneurons orchestrate memory microcircuits. Neuron 112, 201–208.e204 (2024). 10.1016/j.neuron.2023.10.013

15 d’Aquin, S., et al. Compartmentalized dendritic plasticity during associative learning. Science 376, eabf7052 (2022). 10.1126/science.abf7052

16 Topolnik, L. & Tamboli, S. The role of inhibitory circuits in hippocampal memory processing. Nat Rev Neurosci 23, 476–492 (2022). 10.1038/s41583-022-00599-0

17 Tzilivaki, A. et al. Hippocampal GABAergic interneurons and memory. Neuron 111, 3154–3175 (2023). 10.1016/j.neuron.2023.06.016

18 Dunsmoor, J. E., Niv, Y., Daw, N. & Phelps, E. A. Rethinking Extinction. Neuron 88, 47–63 (2015). 10.1016/j.neuron.2015.09.028

19 Bouton, M. E., Maren, S. & McNally, G. P. Behavioral and Neurobiological Mechanisms of Pavlovian and Instrumental Extinction Learning. Physiol Rev 101, 611–681 (2021). 10.1152/physrev.00016.2020

20 Izquierdo, I., Furini, C. R. & Myskiw, J. C. Fear Memory. Physiol Rev 96, 695–750 (2016). 10.1152/physrev.00018.2015

21 Ressler, K. J. et al. Post-traumatic stress disorder: clinical and translational neuroscience from cells to circuits. Nat Rev Neurol 18, 273–288 (2022). 10.1038/s41582-022-00635-8

22 Lebois, L. A. M., Seligowski, A. V., Wolff, J. D., Hill, S. B. & Ressler, K. J. Augmentation of Extinction and Inhibitory Learning in Anxiety and Trauma-Related Disorders. Annu Rev Clin Psychol 15, 257–284 (2019). 10.1146/annurev-clinpsy-050718-095634

23 Vervliet, B., Craske, M. G. & Hermans, D. Fear extinction and relapse: state of the art. Annu Rev Clin Psychol 9, 215–248 (2013). 10.1146/annurev-clinpsy-050212-185542

24 Li, W. G. et al. Input associativity underlies fear memory renewal. Natl Sci Rev 8, nwab004 (2021). 10.1093/nsr/nwab004

25 Gu, X. et al. Dynamic tripartite construct of interregional engram circuits underlies forgetting of extinction memory. Mol Psychiatry 27, 4077–4091 (2022). 10.1038/s41380-022-01684-7

26 Chen, M. B., Jiang, X., Quake, S. R. & Sudhof, T. C. Persistent transcriptional programmes are associated with remote memory. Nature 587, 437–442 (2020). 10.1038/s41586-020-2905-5

27 Sun, W. et al. Spatial transcriptomics reveal neuron-astrocyte synergy in long-term memory. Nature 627, 374–381 (2024). 10.1038/s41586-023-07011-6

28 Peters, J., Dieppa-Perea, L. M., Melendez, L. M. & Quirk, G. J. Induction of fear extinction with hippocampal-infralimbic BDNF. Science 328, 1288–1290 (2010). 10.1126/science.1186909

29 Wang, Q. et al. Fear extinction requires ASIC1a-dependent regulation of hippocampal-prefrontal correlates. Sci Adv 4, eaau3075 (2018). 10.1126/sciadv.aau3075

30 Meyer, H. C. et al. Ventral hippocampus interacts with prelimbic cortex during inhibition of threat response via learned safety in both mice and humans. Proc Natl Acad Sci U S A 116, 26970–26979 (2019). 10.1073/pnas.1910481116

31 Szadzinska, W. et al. Hippocampal Inputs in the Prelimbic Cortex Curb Fear after Extinction. J Neurosci 41, 9129–9140 (2021). 10.1523/jneurosci.0764-20.2021

32 Nguyen, R., Koukoutselos, K., Forro, T. & Ciocchi, S. Fear extinction relies on ventral hippocampal safety codes shaped by the amygdala. Sci Adv 9, eadg4881 (2023). 10.1126/sciadv.adg4881

33 Allen, W. E. et al. Thirst-associated preoptic neurons encode an aversive motivational drive. Science 357, 1149–1155 (2017). 10.1126/science.aan6747

34 Pettit, N. L., Yap, E. L., Greenberg, M. E. & Harvey, C. D. Fos ensembles encode and shape stable spatial maps in the hippocampus. Nature 609, 327–334 (2022). 10.1038/s41586-022-05113-1

35 Fanselow, M. S. & Dong, H. W. Are the dorsal and ventral hippocampus functionally distinct structures? Neuron 65, 7–19 (2010). 10.1016/j.neuron.2009.11.031

36 Marcos, P. & Covenas, R. Regulation of Homeostasis by Neuropeptide Y: Involvement in Food Intake. Curr Med Chem 29, 4026–4049 (2022). 10.2174/0929867328666211213114711

37 Gotzsche, C. R. & Woldbye, D. P. The role of NPY in learning and memory. Neuropeptides 55, 79–89 (2016). 10.1016/j.npep.2015.09.010

38 Tasan, R. O. et al. The role of Neuropeptide Y in fear conditioning and extinction. Neuropeptides 55, 111–126 (2016). 10.1016/j.npep.2015.09.007

39 Michel, M. C. et al. XVI. International Union of Pharmacology recommendations for the nomenclature of neuropeptide Y, peptide YY, and pancreatic polypeptide receptors. Pharmacol Rev 50, 143–150 (1998).

40 Raut, S. B. et al. Diverse therapeutic developments for post-traumatic stress disorder (PTSD) indicate common mechanisms of memory modulation. Pharmacol Ther 239, 108195 (2022). 10.1016/j.pharmthera.2022.108195

41 Wang, H. et al. A tool kit of highly selective and sensitive genetically encoded neuropeptide sensors. Science 382, eabq8173 (2023). 10.1126/science.abq8173

42 Dong, H. W., Swanson, L. W., Chen, L., Fanselow, M. S. & Toga, A. W. Genomic-anatomic evidence for distinct functional domains in hippocampal field CA1. Proc Natl Acad Sci U S A 106, 11794–11799 (2009). 10.1073/pnas.0812608106

43 Wieland, H. A., Willim, K. & Doods, H. N. Receptor binding profiles of NPY analogues and fragments in different tissues and cell lines. Peptides 16, 1389–1394 (1995). 10.1016/0196-9781(95)02028-4

44 Loh, K., Herzog, H. & Shi, Y. C. Regulation of energy homeostasis by the NPY system. Trends Endocrinol Metab 26, 125–135 (2015). 10.1016/j.tem.2015.01.003

45 DeNardo, L. A. et al. Temporal evolution of cortical ensembles promoting remote memory retrieval. Nat Neurosci 22, 460–469 (2019). 10.1038/s41593-018-0318-7

46 Garner, A. R. et al. Generation of a synthetic memory trace. Science 335, 1513–1516 (2012). 10.1126/science.1214985

47 Trouche, S., Sasaki, J. M., Tu, T. & Reijmers, L. G. Fear extinction causes target-specific remodeling of perisomatic inhibitory synapses. Neuron 80, 1054–1065 (2013). 10.1016/j.neuron.2013.07.047

48 Hagihara, K. M. et al. Intercalated amygdala clusters orchestrate a switch in fear state. Nature 594, 403–407 (2021). 10.1038/s41586-021-03593-1

49 Josselyn, S. A. & Frankland, P. W. Memory Allocation: Mechanisms and Function. Annu Rev Neurosci 41, 389–413 (2018). 10.1146/annurev-neuro-080317-061956

50 Hao, Y. et al. Integrated analysis of multimodal single-cell data. Cell 184, 3573–3587 e3529 (2021). 10.1016/j.cell.2021.04.048

51 Finak, G. et al. MAST: a flexible statistical framework for assessing transcriptional changes and characterizing heterogeneity in single-cell RNA sequencing data. Genome Biol 16, 278 (2015). 10.1186/s13059-015-0844-5

